# RecA levels modulate biofilm development in *Acinetobacter baumannii*

**DOI:** 10.1101/809392

**Authors:** Carly Ching, Merlin Brychcy, Brian Nguyen, Paul Muller, Alicyn Reverdy, Margaret Downs, Samuel Regan, Breanna Isley, William Fowle, Yunrong Chai, Veronica G. Godoy

**Author notes:** Corresponding Author: Veronica G. Godoy.

## Abstract

Infections caused by *Acinetobacter baumannii,* a Gram-negative opportunistic pathogen, are difficult to eradicate due to the bacterium’s propensity to quickly gain antibiotic resistances and form biofilms, a protective bacterial multicellular community. The *A. baumannii* DNA damage response (DDR) mediates the antibiotic resistance acquisition and regulates RecA in an atypical fashion; both RecA^Low^ and RecA^High^ cell types are formed in response to DNA damage. The findings of this study demonstrate that the levels of RecA can influence the development, formation and dispersal, of biofilms through the global biofilm regulator *bfmR*. RecA loss results in surface attachment and prominent biofilms, while elevated RecA leads to diminished attachment and dispersal. These findings suggest that the challenge to treat *A. baumannii* infections may be explained by the induction of the DDR, common during infection, as well as the delicate balance between maintaining biofilms in low RecA cells, and promoting mutagenesis and dispersal in high RecA cells. This study underscores the importance of understanding the fundamental biology of bacteria to develop more effective treatments for infections.

## Introduction

*Acinetobacter baumannii* is an emerging Gram-negative opportunistic pathogen and one of the ESKAPE pathogens, a group of bacteria responsible for most hospital-acquired infections(Rice, 2008). *A. baumannii* outbreaks in hospitals are difficult to eradicate, due to increased multi-drug resistance (MDR)(Peleg et al., 2008) and its ability to form biofilms(Eze et al., 2018). *A. baumannii* infections are very dangerous to immunocompromised individuals, causing various illnesses, including pneumonia, septicemia, and wound infections (Smith et al., 2007).

Generally, when gene products involved in antibiotic binding or processing on the chromosome are mutated in a way that impacts their function, resistance can be acquired(Blair et al., 2014). One response pathway underlying multidrug resistance (MDR) is the DNA damage response (DDR). Mutagenesis results from the induction of error-prone DNA polymerase genes, which are part of the DDR regulon(Cirz et al., 2007; Fuchs and Fujii, 2013). We have shown that in *A. baumannii*, RecA-dependent induction of multiple error-prone polymerases in response to DNA damage leads to clinically relevant antibiotic resistance (Norton et al., 2013). In *Escherichia coli* and many other bacteria, the cells’ main recombinase, RecA, and the global transcriptional repressor, LexA, manage the DDR, also known as the SOS response(Little and Mount, 1982). In contrast, there is no known LexA homolog in *A. baumannii*, indicating that the A. baumannii DDR circuitry is regulated differently (Ching et al. 2017; Hare et al. 2014; MacGuire et al. 2014; Norton et al. 2013). Our lab has also shown that in response to DNA damage, there are two RecA cell types, low *recA* expression (RecA^Low^) and high *recA* expression (RecA^High)^, regulated by a 5’untranslated region in the *recA* transcript (Ching et al., 2017; MacGuire et al., 2014).

Bacterial cells within biofilms are often less sensitive to chemical and physical challenges, including antibiotics (Anderl and Franklin, 2000; Singh et al., 2016). Like many biological processes, the biofilm cycle in *A. baumannii* is not fully understood, although key biofilm genes are known, including the biofilm master regulator gene *bfmR* that controls adhesive *csu* pili and genes important for virulence and desiccation tolerance(Farrow et al., 2018; Tomaras et al., 2008, 2003). It is also known that genes encoding Bap protein and efflux pumps are important for biofilms(Brossard and Campagnari, 2012; He et al., 2015; Loehfelm et al., 2008; Tomaras et al., 2008, 2003). The BfmR response regulator is part of a two-component system (BfmRS) that results in BfmR phosphorylation (Farrow et al., 2018; Geisinger and Isberg, 2015; Russo et al., 2016; Tomaras et al., 2008; Wang et al., 2018). There may be additional factors regulating *bfmR* expression and it is unknown whether de-repression by phosphorylated BfmR, with lower affinity for its operator site(Draughn et al., 2018), is sufficient for induction. The *A. baumannii* biofilm extracellular matrix contains poly-ß-(1-6)-*N*-acetyl-glucosamine, mannose and extracellular DNA(Bales et al., 2013; Choi et al., 2009; Hardouin et al., 2014; Sahu et al., 2012).

In brief, the DDR and the biofilm cycle are two crucial pathways that bacteria use for survival under stressful environmental conditions and there is evidence suggesting a link between these two pathways(Linares et al., 2006; Takajashi et al., 1995). The mechanisms linking the DDR and biofilms vary between bacterial species. For example, in *Streptococcus mutans*, *Escherichia coli* (Costa et al., 2014; Inagaki et al., 2009), and *Clostridium difficile* (Walter et al., 2015), there is a direct correlation between DDR and biofilm formation in which both pathways are induced in response to environmental signals. However, we showed that during *Bacillus subtilis* biofilm development, induction of the DDR, measured by levels of oxidative stress and gene expression of a *lexA* -dependent gene (not *recA* or *lexA* directly), shuts off biofilm matrix genes (Gozzi et al., 2017). This suggested an inverse correlation between DDR and biofilm formation in which DDR signaling complements the biofilm cycle; DDR-induced cells could leave the biofilm to search for new niches. Altogether, the varied relationships between DDR and the biofilm cycle are intriguing and show that cells have established different ways to balance these strategies, which could have downstream effects in treatment and clinical practices.

Here, we demonstrate that DDR and the biofilm cycle also intersect in *A. baumannii*. We show that the loss of RecA in *A. baumannii* results in increased cell-to-surface adherence, in part due to elevated expression of *bfmR* with a concomitant increase in Csu pili. We further identify the novel regulation of *bfmR* through the RecA-dependent UmuDAb gene product. UmuDAb is in the same family as LexA and regulates some of the DDR genes in *A. baumannii* (Hare et al., 2014). We observed that DNA – damaged biofilms shut off gene expression of the attachment pili coded by *csuAB* and turn on *recA*, allowing for cell dispersal. The observed inverse relationship between DDR and biofilm formation may lead to a heterogeneous cell population combining physical (biofilm) and genetic (elevated mutagenesis) protection to environmental challenges, including antibiotics.

## Results

### Loss of RecA promotes biofilms in *A. baumannii*

*A. baumannii* ATCC 17978 Δ*recA* cells (*recA*::km, Table S1 (Aranda et al., 2011)) grown in standard LB medium and shaking conditions consistently displayed observable surface attachment on the sides of glass tubes (Fig. 1A, red arrow). In comparison, the parental wild-type strain had little to no surface attachment (Fig. 1A). Intriguingly, this observation suggested that *A. baumannii* Δ*recA* cells form a more prominent biofilm than parental WT cells. Since the lack of RecA can presumably affect bacterial growth (Cox et al., 2008), we first performed growth measurements of free-living planktonic WT and Δ*recA* cells. We also measured growth in Δ*recA* cells complemented with a low-copy plasmid, pNLAC1(Luke et al., 2010), containing *recA* under its own promoter (Ching et al., 2017) (Δ*rec*A (pNLAC1-*recA*), Table S1), which we will refer to as the complemented Δ*recA* mutant or Δ*recA-*c. We observed a slight growth disadvantage in Δ*recA* cells when the starting inoculum size was <10^7^ cells that was rescued by the extrachromosomal copy of *recA* (Fig S1A & B). Our result is largely consistent with a previous report that the Δ*recA* strain showed no noticeable growth defects compared to the parental WT strain(Aranda et al., 2011).

**Figure 1.**
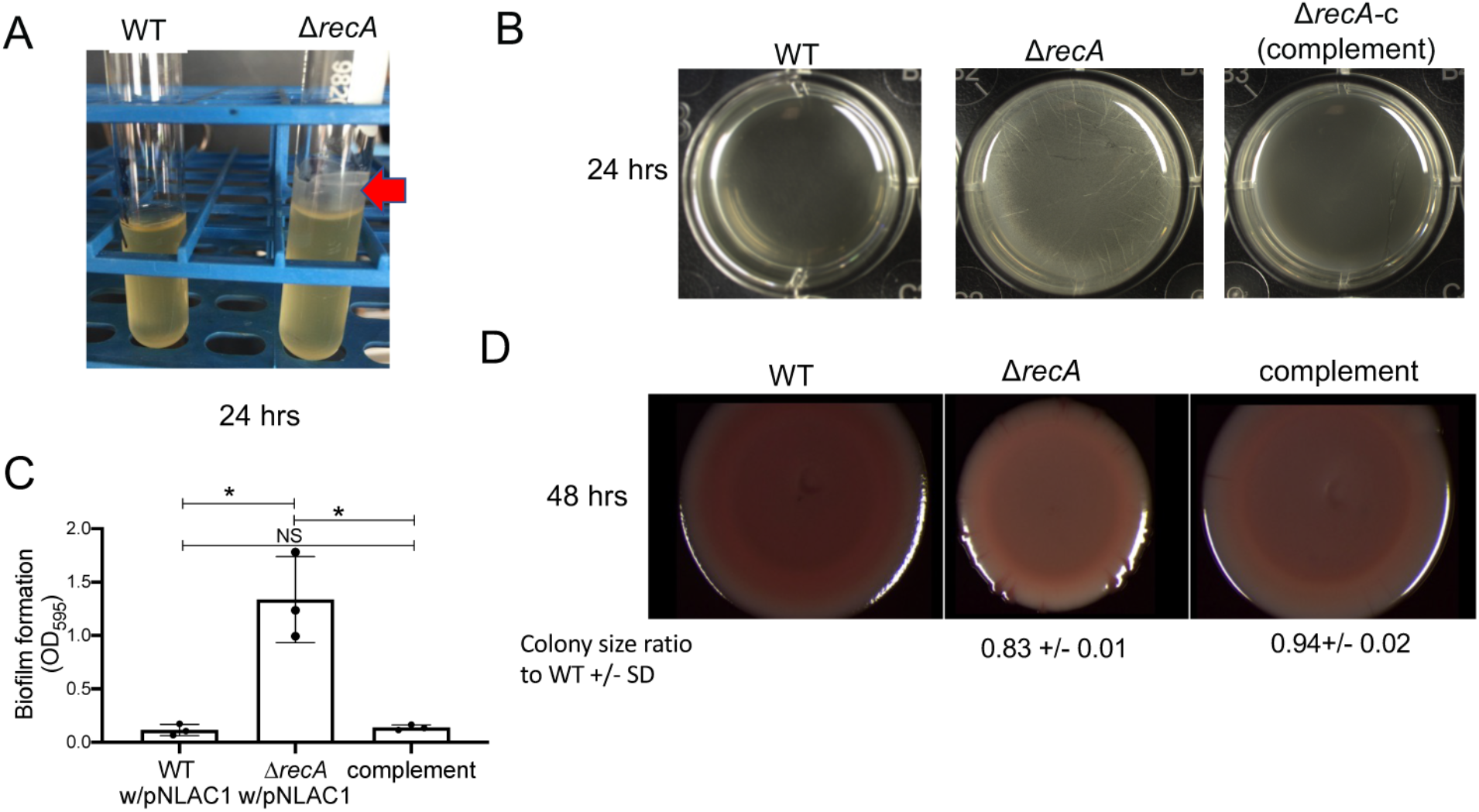
RecA negatively influences surface attachment and biofilm formation. **(A)** Saturated Δ*recA* cell cultures show visible growth on the side surface of glass test tubes (red arrow) compared to WT cells. (**B)** Δ*recA* forms visible pellicles biofilms at the air-liquid interface earlier than its isogenic WT counterpart. Biofilm pellicles of WT (with empty PNLAC1 plasmid), Δ*recA* (with empty PNLAC1 plasmid), and the complemented strain (Δ*recA*(pNLAC1-*recA, ΔrecA-*c) at 24 hrs. Representative images are shown. (**C)** Δ*recA* cells adhere to the abiotic surface better than WT and the Δ*recA*-c strain. Graph shows absorbance at 595 nm for Crystal Violet staining of polystyrene surface-adhered cells at 24 hrs. Experiments were performed in triplicate and error bars shown represent standard deviation of the mean. An unpaired two-tailed T-test was performed for statistical analysis, * =P < 0.05, NS=not significant. Brackets above the columns indicate the comparisons made. (**D**) RecA influences colony biofilm architecture and coloration. Colony biofilms of WT, Δ*recA*, and Δ*recA*-c strains on Congo Red plates at 48 hrs. Numbers underneath the images represent the average diameter of 3 colonies from images taken at the same magnification and settings in 3 independent experiments relative to the WT colony diameter set as 1. The error represents the standard deviation of the mean.

We next assayed pellicle biofilm formation (formed at the air-liquid interface) of Δ*recA* and WT cells, as indicated in materials and methods. WT cells did not form a visible pellicle (Fig. 1B, left) at 24 hrs while Δ*recA* cells (Fig. 1B, middle) formed a visible and opaque pellicle as early as 24 hrs. At 48 and 72 hrs, the WT cells formed a smooth pellicle and with striations while the Δ*recA* pellicle was thicker and granular on the surface (Fig. S2). Cell surface attachement along the sides/bottom of wells of pellicle biofilms of WT and Δ*recA* strains containing the empty plasmid, along with the Δ*recA-*c strain, was measured by Crystal Violet staining. We observed a significant increase in surface-attached cells in Δ*recA* biofilms at 24 hrs compared to WT cells (Fig. 1C). At 24 hrs, pellicle formation of the complemented strain was not significantly different than the parental strain (Fig 1B, right), and the surface architecture was similar to WT (Fig. S2). These data suggest that the loss of RecA leads to a increased biofilm formation.

Colony biofilm formation is indicated by growth at the air-surface interface (Vlamakis et al., 2013). Typically, robust colony biofilms have more wrinkling surface architecture due to differential cell death and mechanical forces due to increased extracellular matrix (Asally et al., 2012). Thus, we grew colony biofilms on solid medium plates containing Congo red and Coomassie brilliant blue, dyes that detect changes in biofilm matrix components (Ghodke et al., 2019; Surgalla and Beesley, 1969). At 48 hrs, the Δ*recA* colony biofilm had differential coloring to that of the WT and complemented strain, indicative of increased biofilm matrix production (Fig 1D). We noticed that the Δ*recA* colony biofilms also had decreased diameter size throughout the length of the experiment. Notably, the complemented Δ*recA* colony is similar in size to WT but with an intermediate coloration (Fig. 1D). We have previously observed partial complementation of colony biofilm phenotypes with plasmid -borne ectopic gene copies, possibly due to the sensitivity of the process to gene dosage (Ching et al., 2018). The data obtained from these experiments suggest that the Δ*recA* strain forms robust colony biofilms.

### The inverse relationship between RecA and biofilm is not exclusive to *A. baumannii* ATCC 17978

To assess whether this finding was exclusive to *A. baumannii* ATCC 17978, we used a transposon derivative of *A. baumannii* AB5075 in which the transposon is inserted approximately in the middle of the RecA open reading frame (ORF) (Gallagher et al., 2015) (Table S1). *A. baumannii* AB5075 is a highly virulent clinical isolate (Jacobs et al., 2014). For pellicle biofilms at 48 hrs, the AB5075 *recA::Tn* mutant had significantly increased cell adherence compared with the parental strain (Fig. S3A). Furthermore, the AB5075 *recA::Tn* colony biofilm is also smaller than the parental strain and displays differences in colony morphology, including changes in opacity and striations on the edge (Fig. S3B). These data suggest that the observation that RecA negatively impacts biofilm development is not exclusive to the ATCC 17978 strain.

### RecA modulates the biofilm matrix in *A. baumannii*

Since Δ*recA* cells form prominent biofilms, we were curious to visually observe differences between Δ*recA* and WT biofilms, especially regarding the extracellular matrix. We thus analyzed Δ*recA* and WT pellicle biofilms by scanning electron microscopy (SEM). To observe pellicle biofilms from the WT and Δ*recA* strains we used a fixing procedure that takes advantage of cationic dyes binding to negatively charged polysaccharides that preserves them for imaging (Erlandsen et al., 2004). We observed noticeable differences between the matrix and biofilm structures of the WT and Δ*recA* strains (Fig. 2A). At 24 hrs, WT cells have some matrix and connections between cells, while Δ*recA* cells were embedded in an observable thick matrix (Fig. 2A). To determine and quantify whether the differences in the biofilm matrix between the WT and Δ*recA* strains were partially due to changes in polysaccharides, total extracellular matrix was extracted from stationary phase cells and carbohydrate content was measured. Δ*recA* cells had a nearly 3-fold increase in total carbohydrates compared the WT cells (Fig. S4). The complemented strain, Δ*recA-*c, had a similar carbohydrate level to WT cells (Fig. S4). It is possible that in addition to increased sugar matrix content we may also be visualizing other phenomena, such as increased pili, which led to further investigation below.

**Figure 2.**
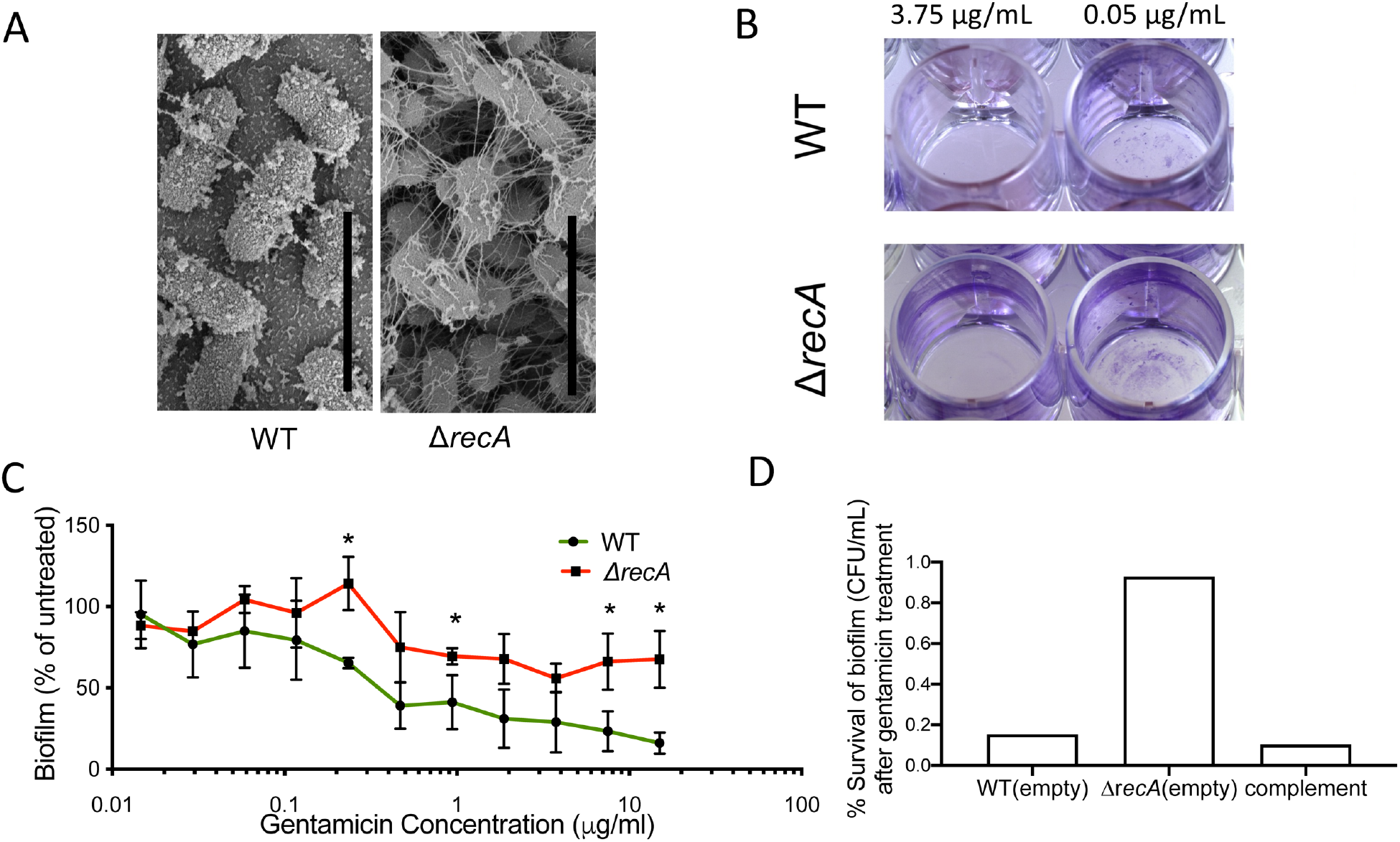
Biofilm formation influences antibiotic challenge. (**A**) Δ*recA* biofilms have increased observable matrix compared to WT biofilms. Scanning electron microscopy (SEM) of WT and Δ*recA* biofilms collected on coverslips at 24 hrs. Scale bar is 3μm. **(B)** Δ*recA* biofilms withstand gentamicin treatment better than WT biofilms. Images of Crystal Violet staining of 72 hrs. mature biofilms formed by WT and Δ*recA* strains after 24 hrs. of gentamicin treatment at 3.75 µg/mL and 0.05 µg/mL. **(C)** Percentage eradication of biofilms (A595 of exposed biofilm/A595 of untreated parental biofilm) treated with a range of gentamicin concentrations for WT and Δ*recA* cells. Experiments were performed in triplicate and error bars represent standard deviation. An unpaired two-tailed T-test was performed to compare % eradication between WT and Δ*recA* strains for each concentration, * = P < 0.05. **(D)** Surface-attached cells with and without gentamicin treatment (15 µg/mL), were deposited on plates and CFUs were counted (representative data from biological replicate with 8 technical replicates each). Graph displays percent survival (CFU/mL) compared to the respective untreated biofilm.

### Δ*recA* surface-attached cells withstand antibiotic treatment better

Biofilms have been noted to protect bacteria from antibiotic exposure (Anderl and Franklin, 2000; Singh et al., 2016). To determine whether there were any differences in the antibiotic susceptibility of the WT and Δ*recA* biofilms compared to planktonic cells, surface-attached WT and Δ*recA* biofilms were treated with the bactericidal aminoglycoside gentamicin. First, we determined the MIC of amicin for exponentially growing planktonic cells in YT medium and found it to be the same for both WT and Δ*recA* cells (1.88 µg/mL, error within 2-fold). Aranda *et al*. determined the MIC of planktonic cells to Amikcin and Tobramycin, both aminoglycosides, and reported that these were also the same for the WT and Δ*recA* strains (1.5 µg/mL and 0.38 µg/mL, respectively) (Aranda et al., 2011).

To assess whether biofilms withstood gentamicin exposure differently to planktonic cells, the spent growth medium from 72 hr pellicle biofilms were removed, the wells were washed to eliminate non-attached cells and gentamicin was added in fresh growth medium at different concentrations, and wells were incubated for 24 hrs. After exposure to gentamicin, the growth medium and non-attached cells were removed, and Crystal Violet staining was performed as before. The data are presented as the percentage of untreated biofilms of each respective strain. For the Δ*recA* strain, the percentage of surface-attached cells remaining after antibiotic treatment plateaus above 50% relative to no treatment after exposure to 0.47–15 μg/mL of gentamicin (Fig. 2C), representing over 3-fold excess of Δ*recA* surface-attached cells compared to WT (at 15 μg/mL the percentage of WT surface-attached cells decreases to ∼16% relative to no treatment). This suggests that that Δ*recA* withstood antibiotic exposure better (Fig. 2B).

To determine if the remaining surface-attached cells were viable, we directly measured the viability of surface-attached cells from pellicle biofilms of WT and Δ*recA* strains containing the empty plasmid, along with the Δ*recA-*c strain, with and without gentamicin treatment, by plating for viable cells (CFUs). We found that with gentamicin exposure (15 ug/mL), the Δr*ecA* strain had ∼ 5-fold increase in percent survival compared to the WT strain, while the complement had similar survival to WT (Fig. 2D).

### Increasing RecA diminishes biofilm formation

Together, our data suggest that there is an inverse relationship between RecA and biofilm development in *A. baumannii.* To further test this relationship, we artificially increased RecA levels in WT cells and expected that a RecA overproducing strain would form poor biofilms with fewer surface-attached cells than WT or Δ*recA* biofilms. Thus, we introduced the plasmid -borne copy of *recA* under its own promoter (pNLAC1-*recA,* Table S1) into the WT strain, which we refer to as *recA*^++^ (Table S1). We first measured the intracellular RecA concentration of *recA*^++^ relative to the WT strain. To do this, we purified *A. baumannii* RecA and *E. coli* RecA to test whether an antibody raised against *E. coli* RecA recognizes *A. baumannii* RecA equally well. There was no difference between the detection of the *E. coli* RecA or *A. baumannii* RecA for the antibody (Fig. S5A&B). Using semi-quantitative Western blotting, we found that the relative level of RecA in *recA*^++^ cells was between 3-to 5-fold higher than WT in both basal and DNA-damage inducing conditions than the parental strain (Fig. S4C). We calculated that after DNA damage, RecA relative levels in the WT strain increased by ∼3-fold relative to basal conditions. Thus, *recA*^++^ at basal conditions has similar levels of RecA as the WT strain treated with 10X MIC of ciprofloxacin (Fig. S5C). The *recA*^++^ strain grown in planktonic conditions had a slight growth defect, similar to Δ*recA* cells (0.15X slower doubling time than WT based on growth curves) (Fig. S1 C&D). We observed that *recA*^++^ biofilms had significantly less surface-attached cells compared to the WT and Δ*recA* biofilms, but still above the baseline reading of a control empty stained well (0.082 +/-0.005) (Fig 3A).

**Figure 3.**
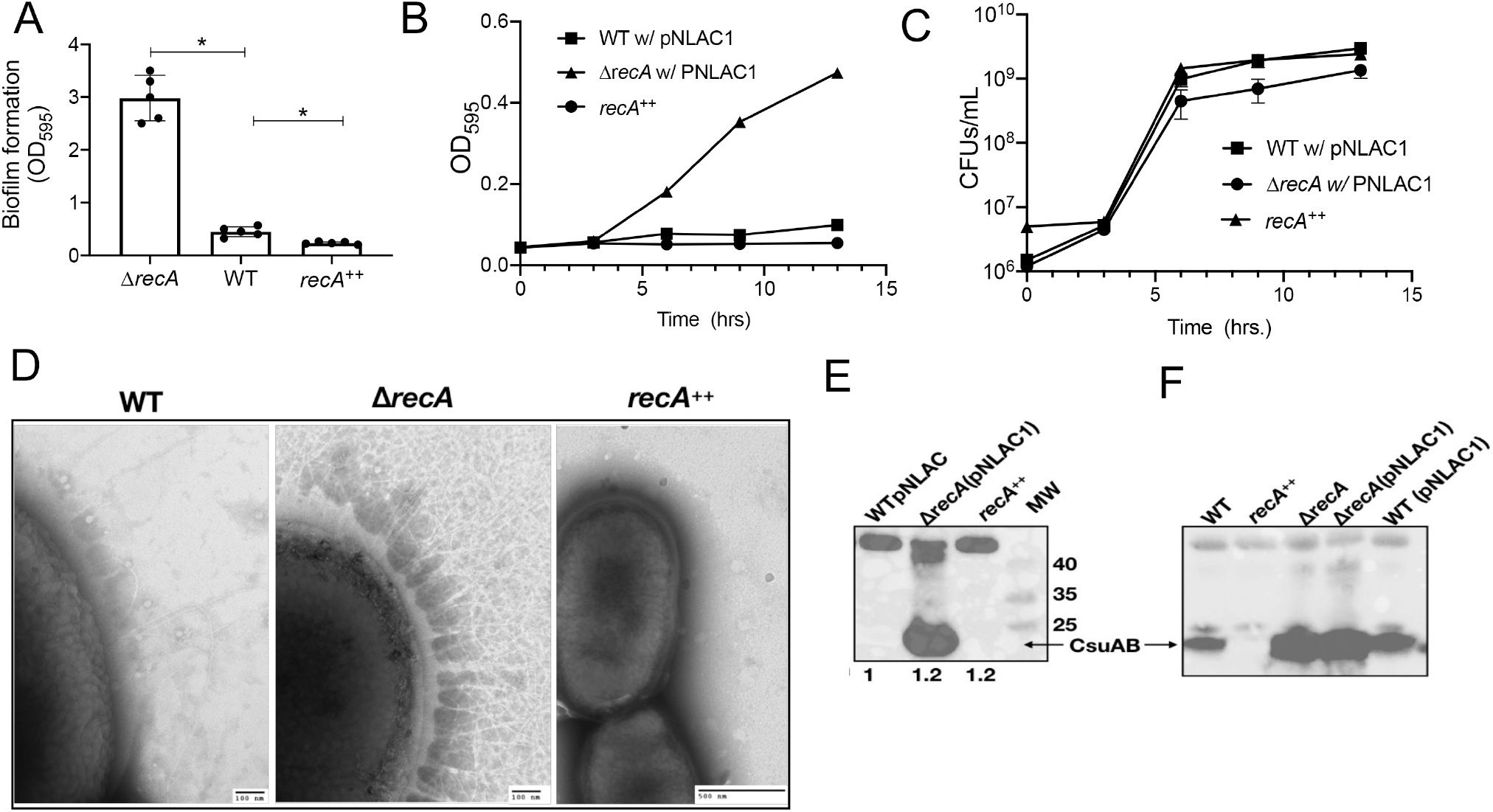
RecA levels modulate *A. baumannii* attachment. (**A**) Cells with high levels of RecA form significantly weaker biofilms than Δ*recA* and WT. Graph displays absorbance at 595 nm for Crystal Violet staining of polystyrene surface-adhered cells from 48 hrs. pellicles of strains Δ*recA,* WT and *recA*^++^ (∼4x more RecA than WT, Fig S5). Error bars represent the standard deviation of the mean. An unpaired two-tailed T-test was performed for statistical analysis. * = P < 0.05. **(B)** Δ*recA* cells attach to surfaces faster than either WT or *recA*^++^. Surface attachment of the isogenic WT, Δ*recA*, and *recA*^++^. All strains contain the pNLAC1 plasmid either empty or carrying the *recA* gene in *recA*^++^. Error bars represent the standard deviation of the mean. (**C**) There is no large difference in growth between strains during the first 13 hrs. of biofilm formation. Colony Forming Units (CFUs)/mL of the respective strains at the same time points as in (B). The error bars represent standard deviation of the mean. Experiment was performed in technical triplicate and biological duplicate. (**D**) The Δ*recA* strain has high density of pili. Transmission electron microscopy images of the pili on the surface of the WT, Δ*recA*, and *recA*^++^ strains. The scale bar for the WT and Δ*recA* strains images is 100 nm and 500 nm for *recA*^++^. Immunoblot for Csu pili isolated from the indicated strains. The total protein ratio of each cell extract from which the pili were isolated compared to WT is shown at the bottom of the image. The >40kD band is non-specific but demonstrates that similar amounts of proteins were loaded per lane. **(F)** Immunoblot for Csu pili from whole cell extracts shows the same effect as in E; there are more Csu pili in Δ*recA*. The empty vector (pNLAC1) does not affect the Csu pili as there is no difference in Csu pili between Δ*recA* cells with or without it. The cell free extracts were loaded at similar concentrations (20 µg of total protein per lane) as also shown by the >40kD band. An anti-Csu polyclonal primary antibody, and an anti-rabbit secondary antibody were used at the dilutions indicated in Materials and Methods for (E) and (F). All lanes are appropriately labelled, and the MW size markers are shown. The CsuAB protein band is highlighted by the arrows.

### RecA levels influence surface attachment

Since planktonic shaking cells for Δ*recA* and *recA^++^*cells is largely consistent with a previous report that the Δ*recA* strain showed no noticeable growth defects compared to the parental WT strain (Aranda et al., 2011) (Fig. S1) in the conditions tested, we concurrently measured biofilm attachment and bacterial growth during the first 13 hours of static growth conditions. We inferred that any differences in biofilms due to growth would be clear at the early stages of biofilm formation while there may be active cell division. To do this, we setup biofilms as indicated in Materials and Methods and measured surface attachment while also determining colony forming units (CFUs) of the whole well combining attached and non-attached cells from 0-13 hrs at regular intervals. Strikingly, at 6 hrs we detected an increase in surface-attached cells for Δ*recA* biofilms compared to WT and *recA*^++^, which both showed little attachment (Fig. 3B). The CFU counts were comparable for the strains (Fig. 3C). These data show that Δ*recA* cells attach to surfaces earlier than WT or the *recA*^++^ strains, as our previous assays suggested (Fig. 1B). This also demonstrates that our findings for biofilm formation are directly comparable and not, in part, due to growth differences. In this experiment, as well as in those one shown previously (Fig. 1B), strains contained the empty plasmid (pNLAC1; Table S1) for direct comparison to the complemented strain. We observed that regardless of pNLAC1 (Fig. 3B, C) Δ*recA* forms robust biofilms, while WT and *recA*^++^ do not. Overall our data indicates an inverse relationship between RecA and biofilm development.

### RecA levels influence the density of attachment pili

Because Δ*recA* cells attached to surfaces faster than WT or *recA*^++^ (Fissg. 3A-C), we wanted to investigate whether Δ*recA* cells had more surface pili to permit attachment. Csu pili are involved in cell adherence in *A. baumannii* biofilms (Gaddy and Actis, 2009; Pakharukova et al., 2018; Rumbo-Feal et al., 2013; Tomaras et al., 2003). To investigate this, we imaged the Δ*recA*, WT and *recA*^++^ strains from saturated cultures using transmission electron microscopy (TEM). Remarkably, Δ*recA* cells displayed a higher density of surface pili than WT or *recA*^++^ cells (Fig. 3D). To determine whether we were observing, in part, Csu pili, we performed a Western blot to detect CsuAB protein from both purified pili (Fig. 3E) and total cell extracts (Fig. 3F) and detected a strong signal for CsuAB pili in Δ*recA* cells.

We next constructed a plasmid containing the promoter of the *csuAB* operon fused to *gfp* (P*_csuAB_*-*gfp*), which was introduced into Δ*recA*, WT, and *recA*^++^ strains. Fluorescence measurements of statically grown cells in YT medium showed that the reporter fluorescence intensity in Δ*recA* was consistently higher compared to WT and *recA*^++^, with *recA*^++^ being consistently the lowest (Fig. 4A). Quantification of fluorescence of single cells (Fig. 4B) shows significantly elevated average fluorescence in Δ*recA* cells compared to *recA*^++^ or WT. Notably, *recA*^++^ fluorescence is significantly lower than WT (Fig. 4B). Our subsequent effort was to quantify the expression levels of *csu* pili genes in biofilm -dwelling cells. Although the qPCR data suggested a higher trend in *csuA* expression in Δ*recA*, the results were deemed statistically insignificant due to the presence of noise. This is likely due to the components in the pellicle samples that may either inhibit PCR, result in poorer mRNA sampling, or due to the inherent heterogeneity of biofilm -dwelling cells (Werner et al., 2004). Nevertheless, in stationary cells, *csuAB* expression is ∼80-fold higher in *A. baumannii* Δ*recA* and lower in *recA^++^* compared to WT cells (Fig. 4C). In summary, our data indicate that Δ*recA* cells have deregulated CsuAB pili.

**Figure 4.**
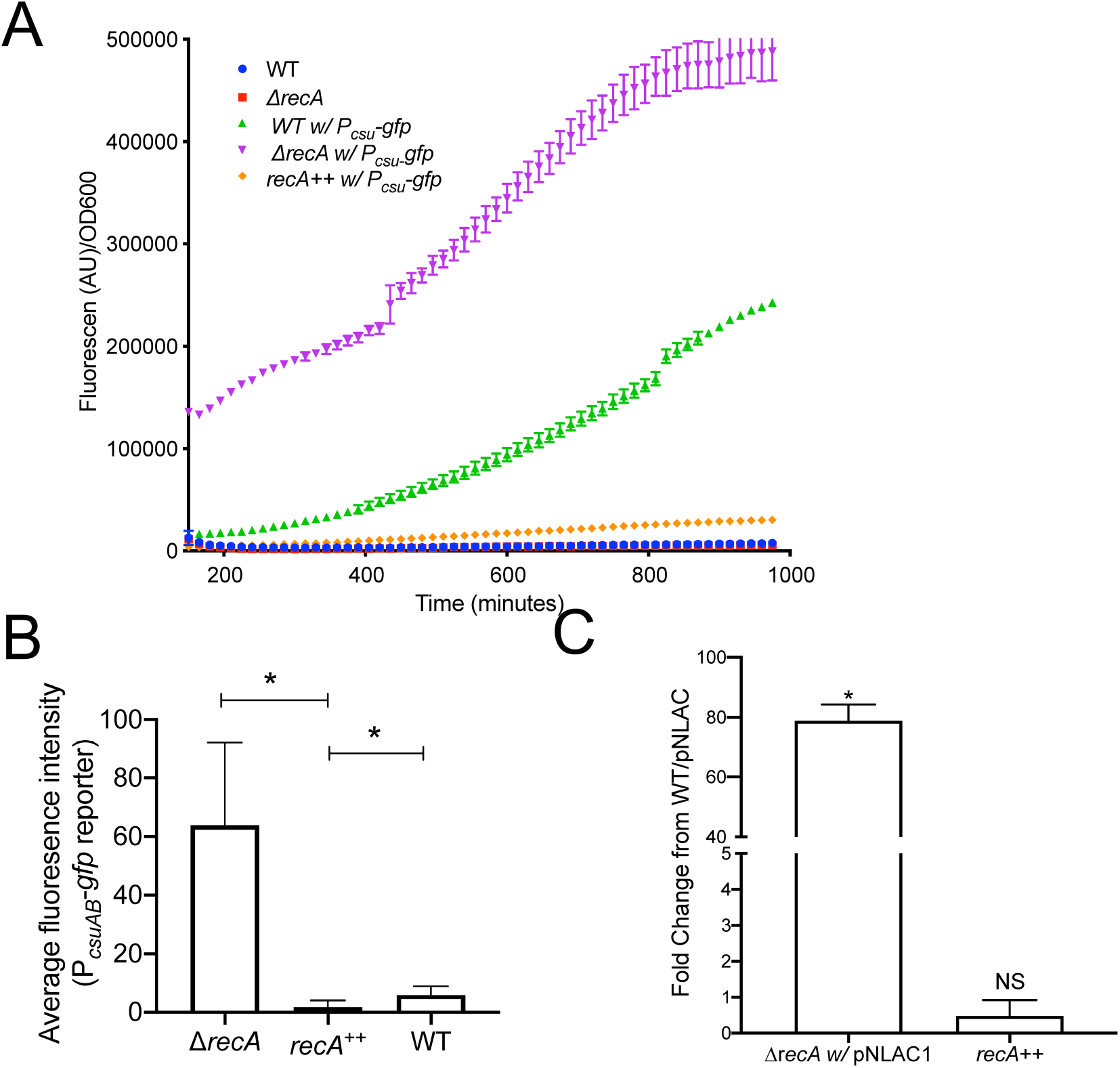
Csu pili expression depends on RecA levels. **(A)** Δ*recA* cells with a Csu fluorescent reporter show elevated and earlier fluorescence compared to WT or *recA*^++^. Fluorescence was measured every 15 mins for a 1000 mins at 25°C in static conditions in a plate reader. (**B**) Quantification of fluorescence of at least 300 single cells demonstrate *csu*-gfp fluorescence dependence on RecA levels; high in Δ*recA*, and low in *recA*^++^. Mean single cell fluorescence with standard deviation is plotted. An unpaired two-tailed T-test was performed for statistical analysis; *P<0.05 (**C**) *csu* transcript is higher in Δ*recA* cells compared to WT and *recA*^++^. qPCR was performed from 24 hrs. saturated cultures in triplicate in three biological replicates to measure *csu* gene expression from the WT, Δ*recA* and *recA*^++^ strains. Expression was standardized to 16S rRNA expression, and the value for the WT strain was set to 1. Error bars represent standard deviation of the mean. An unpaired two-tailed T-Test was used for statistical analysis, compared to gene expression of the WT strain. *= P< 0.05, NS = not significant.

### RecA and *csuAB* expression are inversely correlated in DNA-damaged biofilm pellicles

Due to the absence of a LexA homolog in *A. baumannii*, it is possible that RecA may serve different regulatory roles, as has been previously suggested (Hare et al., 2014). Our results suggested that elevated levels of RecA would lead to decreased CsuAB pili (Figs. 3-4). Thus, we hypothesized that *csuAB* expression would decrease in biofilm cells upon treatment with a DNA damaging agent, which increases RecA. To test this and monitor both RecA and *csuAB* expression, we constructed a plasmid-borne transcriptional reporter of Csu genes (Fig. 4) (P*_csuAB_*-*mCherry*), that was introduced into an *A. baumannii* strain containing a chromosomal P*_recA_*-*gfp* reporter (Ching et al. 2018). 48 hr pellicle biofilms of the double reporter strain were treated with 0.5X MIC of ciprofloxacin (Cip), which induces the DDR but does not kill cells (MacGuire et al., 2014), for an additional 48 hrs. The biofilm pellicles were examined by confocal microscopy to observe differences in gene expression, and in biofilm thickness.

In untreated biofilms (Fig. 5A), most cells had moderate to high expression of the P*_csuAB_*-*mCherry* pili reporter with few expressing P*_recA_*-*gfp* (Fig. 5A). In comparison, cells isolated from Cip-treated biofilms showed dramatic changes (Fig. 5B). Most of the DNA-damaged biofilm cells were fluorescing GFP, an indication of RecA induction, with few cells expressing *mCherry*. There appeared little to no overlap in gene expression between the two reporters, suggestive of mutually exclusive cell types. Moreover, since mCherry and GFP have similar stability (Shaner et al., 2005), they are directly comparable; the gene expression difference is not due to the fluorescence reporters. Most treated cells were also elongated, as cell division inhibition is a common feature of DDR induction (Kreuzer, 2013) that also happens in *A. baumannii*, as we have previously shown (Ching et al., 2017). Moreover, and notably, the untreated pellicles showed high cell density and had a greater thickness (28 µm) than the treated pellicles (22 µm), suggesting that biofilm dispersal occurred in response to DNA damage treatment. It is unlikely that thinning is due to cell death by the Cip treatment since we used sub-MIC levels of Cip. Our data suggest that RecA negatively influences the expression of *Csu* pili genes, necessary for the expression of the Csu pili machinery and biofilm formation.

**Figure 5.**
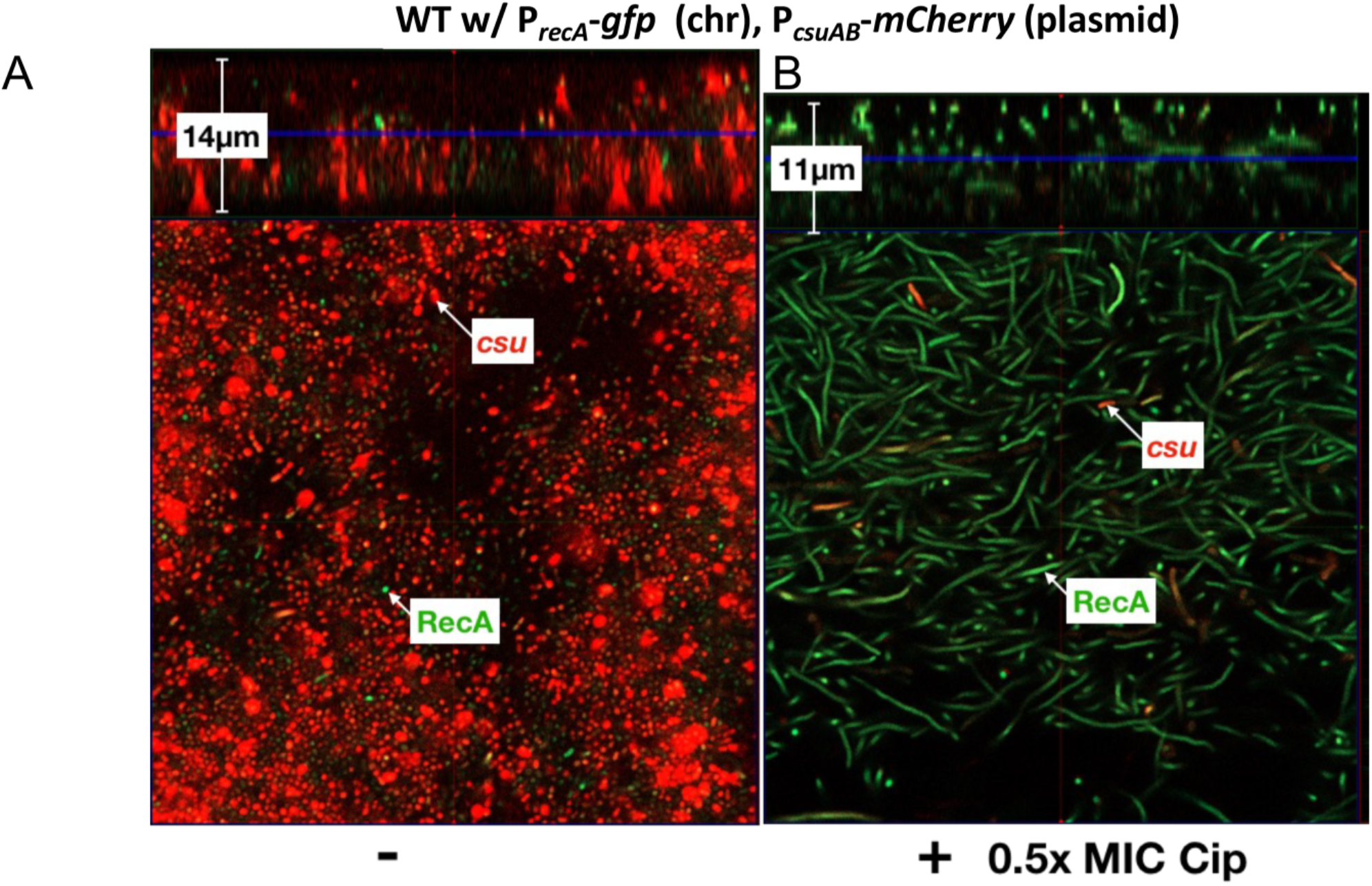
High RecA levels in response to DNA damage led to thinner biofilms. (**A**) Csu pili are highly expressed in biofilm cells. Confocal microscope imaging of a double label *recA*-GFP and *csu*-mCherry reporter strain. The thickness of the biofilm in the mid-Z stack is 14µm as shown in the left-hand side of the figure. There are only a few RecA expressing cells. Csu and RecA cells are labeled in the image. (**B**) RecA levels are high in 48 hrs. ciprofloxacin (0.5X MIC) treated pellicle biofilms. DNA damaged biofilm pellicles images show elongated cells, thinning of the pellicle (mid-Z stack is 11µm) and fewer *csu* expressing cells. Representative images were obtained from a Zeiss Axio Observer.Z1/7 with an excitation/emission of 280/618 for the red channel and of 488/509 for the green channel.

### RecA modulates *bfmR* expression

Csu pili regulation depends on BfmR, a key regulator of biofilm formation in *A. baumannii* (Tomaras et al., 2008). Increased Csu pili in Δ*recA* suggested either higher *bfmR* expression or a more active BfmR. To test whether *bfmR* was differentially expressed in the Δ*recA*, WT, and *recA*^++^ strains we tracked a plasmid borne fluorescent reporter containing the promoter of the *bfmR* operon fused to *mKate2* (P*_bfmR_*-*mKate2*) over time. We found that *bfmR* expression is higher in Δ*recA*, but not in *recA*^++^, compared to WT (Fig. 6A). These results suggest that RecA levels manage *bfmR* expression that in turn controls Csu pili.

**Figure 6.**
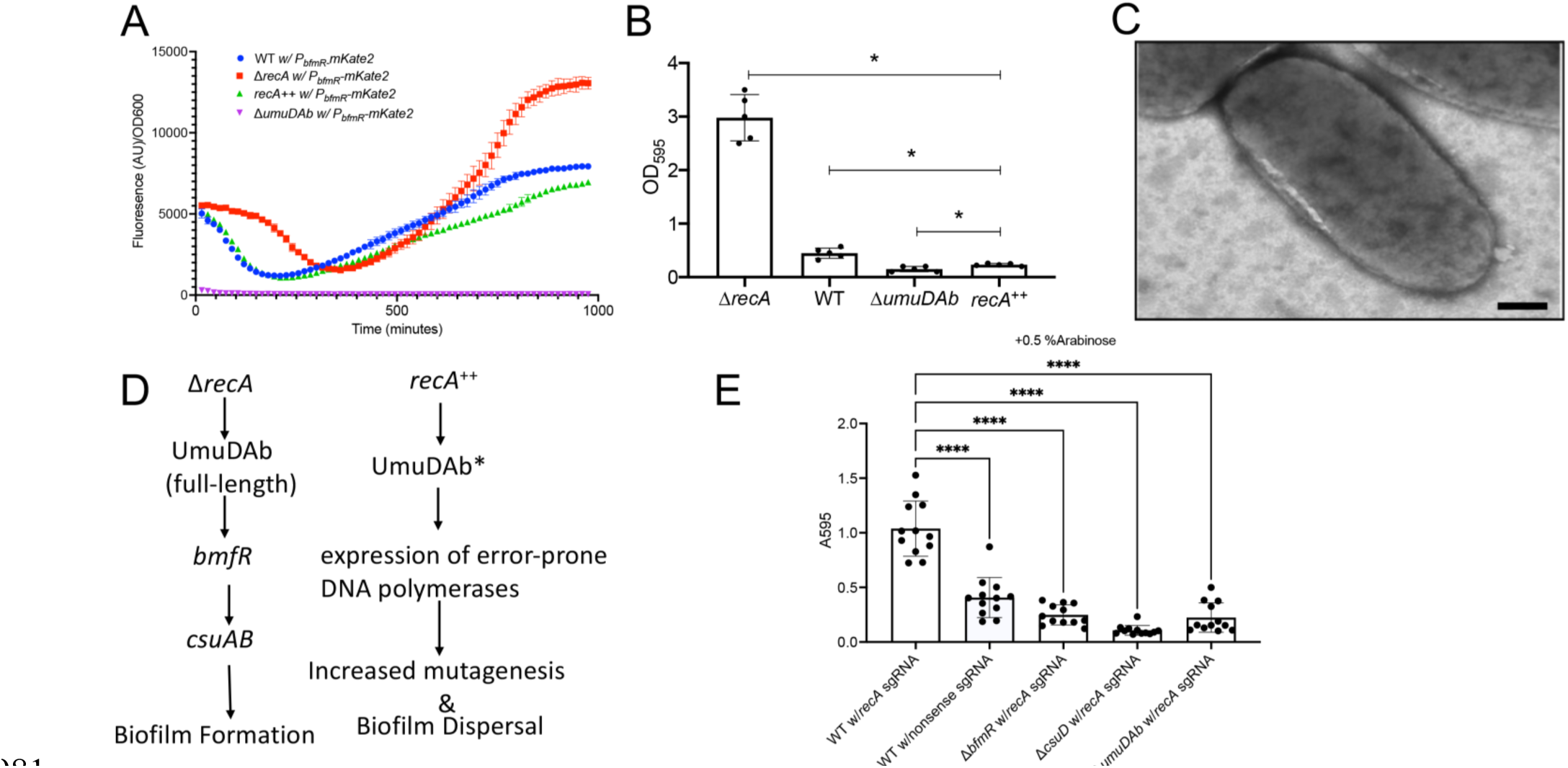
RecA levels modulate biofilm development through *bfmR* and UmuDAb. (**A**) **Δ***recA* cells with a *bfmR* mKate2 fluorescent reporter show earlier fluorescence compared to WT (represented by the blue line), *recA*^++^ (represented by the green line) or Δ*umuDAb* cells (represented by the purple line) while their growth is similar. Fluorescence and OD600 were measured every 15 mins for 1000 mins at 25°C in shaking conditions in a plate reader. (**B**) A strain lacking *umuDAb* form poor biofilms compared to other strains. The Δ*recA* strain forms biofilm significantly better than any of the other strains tested in pairwise statistical analysis shown by the straight line on top of the columns. The WT strain is significantly different to Δ*umuDAb* and *recA*^++^, while Δ*umuDAb* is significantly different to *recA*^++^. An unpaired two-tailed T-test was performed for statistical analysis; *p<0.05. (**C**) Transmission electron micrograph of the Δ*umuDAb* strain shows no evident surface pili. Representative image shown. Scale bar is 100nm. (**D**) Model based on the presented data. Arrows represent positive effect. **(E)** A plasmid-based arabinose inducible CRISPRi system was used to knock down *recA* in WT, Δ*umuDAb,* Δ*bfmR* and Δ*csuD* cells. Graph shows absorbance at 595 nm for Crystal Violet staining of polystyrene respective surface-adhered cells at 48 hrs. Experiments were performed in triplicate and error bars shown represent standard deviation of the mean. An unpaired two-tailed T-test was performed for statistical analysis, **** =P < 0.05.

### UmuDAb, a RecA-dependent transcription factor, regulates BfmR

While *A. baumannii* lacks LexA, UmuDAb (A1S_1389), is a known repressor of some, but not all, DDR genes(Hare et al., 2014). Notably, UmuDAb has a typical helix-turn-helix DNA binding motif at the amino-terminal end that is cleaved by the activated RecA nucleoprotein filament, RecA*, in a way believed to be to that of LexA and other proteins within the same family (Draughn et al., 2018; Hare et al., 2013). Cleavage of UmuDAb by RecA* provides a plausible mechanistic explanation for the observations that high RecA levels (i.e. low full-length UmuDAb) lead to biofilm dispersal. Thus, we hypothesized that UmuDAb may serve as a RecA-dependent regulator of biofilm development.

We constructed an Δ*umuDAb* strain as previously done (Norton et al., 2013) and tested for biofilm formation. We found that the Δ*umuDAb* strain formed significantly weaker biofilms than any other strain previously studied in pairwise comparisons (Fig 6B). In fact, Δ*umuDAb* formed significantly less biofilm than *recA*^++^ (Fig. 6B). Consistent with this observation, TEM showed no visible pili on the surface of Δ*umuDAb* cells (Fig 6C). Furthermore, we detected almost no *bfmR* expression in Δ*umuDAb bfmR* reporter cells compared to WT (Fig 6A). Contrary to the known UmuDAb repressor function(Hare et al., 2014; Witkowski et al., 2016), our findings suggest that UmuDAb has a positive effect on *bfmR* and consequently on Csu pili. Indeed, we have biochemical evidence that full-length UmuDAb binds to the *bfmrR* promoter, while the cleaved version does not (Fig. S6A). Our group is currently further investigating the mechanistic details by which UmuDAb performs this function. Overall, our data suggest a genetic model in which *A. baumannii* biofilm development depends on RecA, UmuDAb, BfmR, and CsuAB (Fig. 6D, expanded upon in discussion). To test this genetically, we designed a plasmid-based arabinose -inducible CRISPRi system and knocked down *recA* in WT, Δ*umuDAb,* Δ*bfmR* and Δ*csuD*(Moon et al., 2017) cells (Fig. S6B). We then compared the pellicle biofilm formation between these strains. As expected, when *recA* was knocked down in WT cells, there was a significant increase in biofilm compared to WT cells with a nonsense single guide RNA (Fig. 6E). Furthermore, and as expected, knocking down RecA in each of Δ*umuDAb,* Δ*bfmR* and Δ*csuD* strains, did not increase biofilms (Fig 6E).

## Discussion

Bacteria largely exist in the environment as biofilms (Costerton et al., 1995). Generally, cells in biofilms have different characteristics and gene expression profiles from their planktonic counterparts. Here, we find that in *A. baumannii,* levels of RecA, a key DDR gene product, influences biofilm development in both ATCC 17978 (Figs. 1-6) or AB5075 (Fig. S3) strains. Our data also show that cells lacking *recA* produce robust biofilms (Fig. 1). Importantly, Δ*recA* biofilms on an abiotic surface expected to be more resilient (Fig. 2) and a higher fraction of cells remain viable after antibiotic treatment (Table 1) compared to the parental strain, providing additional evidence to the concept that biofilms protect cells from antibiotic exposure (Anderl and Franklin, 2000; Singh et al., 2016). Thus, recurring *A. baumannii* infections may be the product of cells remaining adhered to equipment surfaces or implanted devices.

We found that RecA levels influence biofilm formation through Csu pili. Δ*recA* cell adherence depends on higher *csuAB* and increased Csu pili compared to either the WT or the *recA*^++^ strains (Figs. 3, 4). Our results further suggest that *csuAB* expression is due to the induction of *bfmR* (Fig. 6). It has been previously shown that *bfmR* and *csuAB* expression are higher in biofilm cells (Rumbo-Feal et al., 2013), consistent with our findings.

Aranda *et al*. phenotypically characterized the *ΔrecA* insertional mutant used in this study. Free-living *ΔrecA* cells had decreased survival during heat shock, desiccation, UV, and certain antibiotic treatment(Aranda et al., 2011). Importantly, the *ΔrecA* strain had much lower pathogenicity in a mouse model, indicating its importance in this process (Aranda et al., 2011). These results demonstrate the significance of RecA in survival and virulence in planktonic cells. However, our findings demonstrate the importance of investigating both biofilm and free-living states. For example, it has been shown that strong *A. baumannii* biofilm formers are less frequently antibiotic-resistant, consistent with having lower levels of RecA and subsequently mutagenesis (Wang et al., 2018). Additionally, planktonic cells more readily gain higher level antibiotic resistance compared to biofilm cells during exposure to ciprofloxacin (Santos-Lopez et al., 2019). Moreover, biofilm cells were more eradication-resistant (Wang et al. 2018), again consistent with the observation that Δ*recA* surface-attached cells are more difficult to eliminate with antibiotics (Fig. 2, and Table 1). This could be because certain antibiotics have lowered penetration of biofilms (Anderl and Franklin, 2000; Singh et al., 2016) and there is a nutrient gradient within the biofilm that leads to different metabolic states of biofilm cells (Werner et al., 2004). Other explanations include an increase in antibiotic efflux in the biofilm conditions or that the surface of the bacterium is very different in biofilms, and the antibiotic does not interact as efficiently.

Our findings have significant implications for understanding *A. baumannii* survival to antibiotic treatment and possibly antibiotic resistance. We show that upon DNA damage, the DNA damage response is induced, leading to the induction of *recA* with simultaneous shutoff of biofilm genes, leading to thinner biofilms (Fig. 5). We have observed that this occurs through lowered *bfmR* and *csuAB* expression upon elevated RecA levels. Quantification of RecA upon DNA damage, and in *recA^++^*cells, demonstrates how sensitive the biofilm gene expression is to a change in RecA levels (Figs. S4, 1-3). This may be due to UmuDAb sensitivity to cleavage by RecA* (Hare et al., 2013). Our evidence suggests that full-length UmuDAb is an inducer of *bfmR* and of biofilm formation (Fig. 6 and supplementary Fig. S6B). A summary and model of genes, supported by our data, is shown in Fig. 6D. Based on previous findings, we have observed two of RecA cell types, high and low, in planktonic cells (Ching et al., 2017; MacGuire et al., 2014), and there may also be heterogeneity in RecA levels among cells in biofilms. Thus, within a biofilm, the RecA^Low^ cells maintain biofilms with low mutagenic potential, while RecA^High^ cells with high mutagenic potential (Norton et al., 2013) can disperse and search for new niches. This observed inverse relationship can allow the population to combine physical (biofilm) and genetic (elevated mutagenesis) protection to environmental challenges, including antibiotic exposure in a host.

Overall, we have identified an inverse relationship between the DDR and biofilm development in *A. baumannii.* While a similar anti-correlation was observed in *B. subtilis*, *A. baumannii* has a unique DDR that it is not a canonical LexA-dependent SOS system. Thus, our findings provide further insights on LexA-independent DDRs. Here, we uncover different underlying regulatory pathways and genes mediating this inverse relationship. Specifically, the *Gozzi et al* paper does not test RecA directly, and the mechanism is through the accumulation of reactive oxygen species, which induce the canonical DDR and genes known to be in the LexA -dependent regulon(Gozzi et al., 2017). We show that different levels of RecA govern biofilm genes and pili in a LexA-independent system. The direct relationship between RecA and pili has not been reported and notably we have expanded on the RecA-dependent regulon in *A. baumannii*. Although there may be by convergent evolution of the same approach (biofilm and DNA damage responses have mutual exclusivity), we reveal a fundamentally different approach in distantly related bacteria (Gram-negative vs Gram-positive, nonpathogenic vs pathogen).

These results demonstrate the complexity of treating pathogens with a DDR that do not follow the paradigm, such as *A. baumannii* (Ching et al., 2017; MacGuire et al., 2014). Previous work has highlighted the potential for clinical RecA inhibitors to potentiate the effect of antibiotics while hindering antibiotic resistance acquisition by reducing the mutagenic capacity in bacteria (Alam et al., 2016). This indeed may be promising for treating certain bacteria, as demonstrated in *E. coli* (Alam et al., 2016). However, our results show that for *A. baumannii*, inhibiting RecA may lead to robust biofilm formation. It is thus important to understand the relationships between survival strategies in bacteria, and that different bacteria may have different responses to treatment, based on their fundamental biology. Overall, our results show the complex ways that *A. baumannii* robustly survives stress by balancing alternative survival strategies through overlapping genetic pathways.

## Materials and Methods

### Strains and Growth Conditions

The strains and plasmids used are listed in Table S1. *A. baumannii* strains harboring the PNLAC1-*recA* plasmid were constructed as before (Ching et al., 2017). The Δ*umuDAb* strain was constructed using SOE PCR to insert a kanamycin gene cassette in the *umuDAb* gene as before (Norton et al., 2013). The oligonucleotides used are in Table S1. All bacterial cultures were routinely grown in LB medium, unless otherwise noted, and incubated at 37°C with shaking at 225 rpm for liquid cultures. YT medium used is composed of 2.5-g NaCl, 10g Tryptone and 1g Yeast extract per liter. The solid medium contained 1.5% agar (Fisher Bio-Reagents). Antibiotics were used at the following concentrations: Kanamycin (Kan; 35 µg/mL), Gentamicin (Gm; 10 µg/mL), Tetracycline (Tet; 12 µg/mL), and Carbenicillin (Carb; 100 µg/mL).

### Growth curve measurements

Strains were diluted in the growth medium as indicated in the respective figure legends and grown in YT medium in 96-well dishes. All strains were tested at least in triplicate. They were incubated in a plate reader (Biotek Synergy H1) at 37°C with shaking for 24 hrs with measurements of OD_600_ every 15 min.

### Pellicles & Colony Biofilm Formation

To form pellicle biofilms, cells from a saturated culture were inoculated at a 1:1000 dilution into YT liquid medium in either 6, 12 or 24-well non-tissue culture treated (lacking coating which change surface properties) polystyrene plates. The same number of cells, adjusted based on optical density, was added to each well. Plates were then incubated statically at 25°C. A standard Crystal violet staining procedure was used to quantify adherence to the polystyrene surface (Chen et al., 2015). Early biofilms are formed after 24 hrsh of static growth, and between 48-72 hrs. the biofilm has matured. After 72 hrs., the biofilms start to deteriorate. In the time course and gentamicin viable cell count experiments, to correlate viable cells and biofilm formation, samples were taken for CFU counting from 3 wells per experiment. The well content was thoroughly mixed before taking the samples, diluted with PBS, and deposited on LB agar plates. After CFU sampling, the well content was aspirated, washed 3 times with PBS, and stained with Crystal Violet as before (Ching et al., 2018). For colony biofilms, 3 µL of a saturated bacterial culture were spotted on to YT agar plates containing 200µg/mL Congo red and 100µg/mL Coomassie brilliant blue (Congo red plates) (Ching et al., 2018) and incubated for 48 hrs., early colony biofilms, to 96 hrs. at 25°C. Images of biofilms were taken with a Leica MSV269 dissecting scope and a Leica DMC2900 camera, using the same settings. The radius of the biofilm colonies was measured in triplicate from the dissecting microscope images at the same magnification.

For AB5075 colony, R2 medium with 1.5% agar (Reasoner and Geldreich, 1985) was used instead of YT.

To prepare biofilms in which *recA* expression was knockdown by CRISPRi (Brychcy et al., n.d.), 0.5 % L-Arabinose was added to the YT medium to induce expression of the *dCas9* gene and inhition of *recA* expression.

### Scanning Electron (SEM) and Transmission Electron (TEM) microscopy

Biofilms for SEM were prepared by collecting pellicle biofilms on a glass coverslip treated with a solution of 0.1 mg/mL of polylysine (Fisher Scientific). The coverslips with the biofilms were deposited for at least 24 hrs. at 4°C in wells with fixative 1 composed of 25% Glutaraldehyde, 0.1 M Na-Cacodylate buffer (pH 7.2), 0.15% Alcian Blue and 0.15% Safranin. Additional treatment of the coverslips and observation of the biofilms was performed at the Northeastern University Electron Microscopy Core Facility.

For TEM, cultures were grown to saturation in YT liquid medium and streaked for single colonies on YT plates incubated at 37°C. A colony was picked and gently resuspended in 100 µL of phosphate buffered saline (PBS). 10 µL of the resuspended colony was pipetted onto a copper grid, excess liquid wicked away with filter paper, and the grid was dried at room temperature. To negatively stain the samples and observe pili, 10 µL of a solution of 1.5% phosphotungstic acid (PTA) was pipetted onto the cell-containing grids. Liquid excess was wicked away with filter paper, and grids were left to air dry at room temperature. Microscopy was performed with a JEOL JEM-1010 (JEOL USA, Peabody, MA) microscope equipped with a 2k x 2k AMT CCD camera (Advanced Microscopy Techniques, Woburn, MA) at the Northeastern University Electron Microscopy Core Facility.

### Sugar Extraction Assay

The total extracellular matrix was extracted and total carbohydrate content was quantified using a modified protocol from Jiao et al. 2010 (Jiao et al., 2010). Briefly, 500 µL of YT medium saturated cultures of the *A. baumannii* strains were centrifuged at 13,000 rpm for 1 min in pre-weighed microcentrifuge tubes and washed twice with 1X PBS; supernatant was discarded. Cell pellets were dried at 95°C and reweighed. This process was repeated until all dried pellets were approximately equal in weight (differences no more than 20% from each other). Pellets were reconstituted in 100 µL of 1X PBS and 400 µL of a 0.1 M sulfuric acid solution was added to each tube. Cell suspensions were mixed in a vortex every 10 minutes for 1 h.. Then, 250 µL of concentrated sulfuric acid (95% minimum) was added, followed by 50 µL of freshly made 10% phenol solution in water. This mixture was incubated at 95°C for 5 minutes. The absorbance at 492 nm of each sample was determined in a Biotek plate reader (Synergy HT). This measurement represents the total carbohydrates. The absorbance per gram of dry cell was normalized to the values obtained with the parental strain for each 3 biological replicates. An unpaired two-tailed T-Test was used for statistical analysis relative to the parental strain: * = P < 0.005.

### Quantitative measurement of gene expression

To measure *csu* transcripts in WT, Δ*recA*, and *recA*^++^, we took 500µL of 24 hrs. saturated cells. These were harvested by centrifugation and resuspended in 1 mL of RNA Protect Bacteria Reagent (Qiagen) with 30µL of 20 mg/mL lysozyme, incubated at room temperature for 15 min, spun down at 15,000g for 5 min, and cell pellets were stored at -20 °C. Total RNA extraction was carried out by using the Zymo Direct-zol RNA Kit (Zymo) according to manufacturer’s instructions. A NanoDrop One (ThermoFisher) was used to measure the RNA concentration and purity. RNA was converted to cDNA using a high -capacity cDNA reverse transcription kit according to the manufacturer’s protocol (Applied Biosystems). Expression of *csuAB* pili gene expression by RT-qPCR was performed with cDNA using the qPCR primer pairs listed in Table S1 and following the protocol for Fast SYBR Green Master Mix (Applied Biosystems) using a StepOnePlus real-time PCR instrument (Applied Biosystems). The relative gene expression was standardized using endogenous 16S ribosomal RNA expression, and the comparative threshold cycle (ΔΔCT) was calculated for each sample and compared relative to WT expression. Each sample was run in biological and technical triplicates. An unpaired two -tail T test was used for statistical analysis relative to the parental strain (*=P < 0.05).

### Pilus purification

Pilus shear preparations were performed as previously published (Moon et al., 2017). Briefly, a similar number of cells grown on a YT agar plate (OD_600_∼20) were collected in 1.5 mL of 1X PBS, placed on ice for 10 min, and vortexed for 1 min. After centrifugation at 13,000 x g at 4°C, cell supernatants (containing the pili) and the respective pellets (containing pili-less cells) were collected. The pili within the supernatants were precipitated for 20 h with trichloroacetic acid (TCA) (final concentration, 25%), at -20°C. The precipitated pili were collected by centrifugation at 16,000 x g for 10 min at 4°C and resuspended in 50 µL 1X PBS. Total protein was extracted from pili-free cell pellets and whole cell lysates using Bugbuster reagent (Novagen) following the manufacturer’s instructions and quantified using a Bradford assay.

### Purification of His-tagged *A. baumannii* RecA and Immunoblots

The *recA* gene from *A. baumannii* was amplified using the primers RecALICF and RecALICR (Table S1), ligated into the pET-His6-TEV-LIC vector (plasmid 29653; Addgene, Cambridge, MA, USA) using the Ligation Independent Cloning Protocol (Gradia et al., 2017), and introduced by transformation into DH5α *E. coli* cells for plasmid maintenance. The induction of expression and purification methods are as previously described for RecA purification from *E. coli* (Tashjian et al., 2017). RecA purification was confirmed by SDS-PAGE.

For the semi-quantitative western blots, saturated cultures were diluted 1:100 and grown to an exponential phase. Cells were then treated with ciprofloxacin (Cip) at 10X the MIC for 3 h. (Ching et al., 2017). A parallel culture for each strain was left untreated. Cells were collected by centrifugation, and whole-protein lysate was extracted using Bugbuster reagent (Novagen) and Pierce Universal Nuclease according to manufacturer’s instructions. Cell -free lysates were quantified with a Bradford assay following the manufacturer’s instructions (BioRad). Lysate was diluted to 1X with Laemmli buffer and heated for 10 min at 95°C. The samples were separated on a 12% Bis-Tris gel (Invitrogen) in MOPS buffer and transferred to a nitrocellulose membrane (Cafarelli et al., 2013). The blot was developed as before (Cafarelli et al., 2013) using a primary anti-RecA antibody (Abcam) at a 1:10,000 dilution (Norton et al., 2013) and an HRP-labelled secondary goat anti-rabbit antibody (Abcam) diluted at 1:40,000. Image J was used to quantify the density of the RecA bands (Norton et al., 2013). The RecA signal from all samples were compared to a standard of purified RecA protein from *A. baumannii*. Cmmassie Blue staining was used to estimate similar loading of lanes of a parallel gel. The procedure was done in 4 biological replicates. RecA determinations were similar every time.

To measure the relative amount of Csu pili in the different strains, we performed immunoblots with an anti-Csu antibody (gracious gift from Luis Actis). The pili content of each strain was standardized to the total protein measured in the cell-free extracts from which the pili were isolated. To detect Csu pili from whole cell lysates, a similar number of cells were lysed with Bugbuster and Pierce Universal Nuclease as indicated above. The samples were separated in a 12% polyacrylamide gel in MES buffer at 150V for 1 h.. Immunoblot procedures were followed as described above using a 1:1000 dilution of the anti-Csu primary antiserum and a 1:40,000 dilution of the HRP labeled secondary anti-rabbit antibody (Abcam). A ChemiDoc MP imaging system (BioRad) was used for chemiluminescence signal detection from RecA and Csu blots.

### EMSA for UmuDAb to the promoter of *bfmR*

The full-length *umuDAb* and truncated *umuDAb* (*umuD’Ab,* C-terminal end based on previously determined cleavage site (Hare et al., 2013)) genes were tagged with a hexahistidine tag on the C-terminal end and cloned into pET11T (Nguyen et al., 1993). The constructs were introduced into BL21 AI cells and His-UmuDAb and His-UmuD’Ab were overexpressed as previously described (Cafarelli et al., 2013; Studier, 2005). Both proteins were then purified by immobilized metal affinity chromatography as described previously (Cafarelli et al., 2014). PCR amplification with primers (bfmR_umuD_motif_F(Cy3) and bfmR_umuD_motif_R, Table S1) labeled with a Cy3 fluorophore on the 5’ end was used to create the *bfmR* DNA probe. DNA (25 nM) and protein were mixed with binding buffer (10mM Tris-HCl [pH 8], 10mM HEPES, 50mM KCl, 1mM EDTA, 100 ng of salmon sperm DNA, 0.1 mg of bovine serum albumin per ml). Binding reactions were incubated for one h at 37°C. Reaction mixtures (20 ul) were loaded into a 5% nondenaturing Tris-borate-EDTA (TBE) polyacrylamide gel. Binding complexes were separated at 150V for one hour. Cy3-labelled DNA-protein complexes were visualized using a Bio-rad Chemidoc imager. All the EMSAs were repeated thrice.

### Construction of the P*_csu_*-*gfp*, P*_csuAB_*-*mCherry* and P*bfmr-mkate2* expression reporter plasmids and expression assays

*A. baumannii* ATCC 17978 containing a *csu* operon (P*_csuAB_-gfp*) transcriptional reporter on the Carbenicillin (Carb) resistant plasmid pMB1-A(Brychcy et al., n.d.), a modified version of the EcoFlex kit (Moore et al., 2016) containing replication elements necessary for *A. baumannii*, was used to assess the expression of Csu in the WT, *recA*^++^, and Δ*recA* strains. The promoter region of the *csu* operon and of *bfmr* were amplified using the primers csu_f and csu_r, and pBfmR_f and pBfmR_r respectively. The plasmid pBP_lacZ (Table S1; Addgene accession number 72948) and the purified PCR fragment were digested using SphI and NdeI, and the pBP-lacZ plasmid was dephosphorylated using QuickCIP (all enzymes from NEB). The ligation was carried out with T4 DNA ligase (Thermo Fisher Scientific). Colonies were screened using blue-white color in plates with X-Gal and verified by colony PCR and sequencing using Primers pBP_pTU_f and pBP_r. This generated level 0 MoClo plasmids for the promoter elements.

Plasmids with the desired inserts were then used in a combined cloning and ligation reaction, by using 20 fmol of each entry vector (pBP_*pcsu* or pBP_*pbfmr*, pBP_B0012, pBP_e*gfp;* pBP_*mcherry* or pBP_*mkate2*, pBP_pET-RBS and pTU1-A-RFP-AB) combined with 2 µL of Cutsmart, 0.5 µL of BsaI-HFv2, 0.5 µL of T4 DNA Ligase and 1 µL of 10 mM ATP (all reagents from New England Biolabs, Ipswich, MA., except T4 DNA Ligase from Thermo Fisher Scientific). The reaction was cycled 50 times in a thermocycler with a 37 °C digestion step for 2 minutes and a 16 °C ligation step for 5 minutes. Plasmids with no insert were eliminated by digestion with BsaI at 37 °C for 1 h. The enzymes were heat-inactivated at 80°C for 10 min. 5 µL of the ligation mix was introduced by transformation into chemically competent *E. coli* DH5α. White colonies contain the desired insert due to the exchange of the mCherry gene originally in the pMB1-A cloning site and confirmed by colony PCR and sequencing the insert using pBP_pTU_f and pTU_r. Plasmids were introduced by transformation into *A. baumannii* using standard electroporation protocols (MacGuire et al., 2014).

To quantify fluorescence in single cells, at least 300 single cells were counted, and their fluorescence was measured with MicrobeJ from microscope images taken with the same settings(Ducret et al., 2016).

### Construction of *recA* knockdown CRISPRi plasmids

CRISPRi constructs were created according to Brychcy et al.(Brychcy et al., n.d.). Briefly, 5 uL of 100uM of each sgRNA oligonucleotide (recA_ORF_sgRNA_f and recA_ORF_sgRNA_r or nonsense_sgRNA_f and nonsense_sgRNA_r) were annealed in T4 DNA ligase buffer by incubating at 95 °C for 5 minutes and subsequent cooling to 22 °C for 20 minutes. 1 uL of the annealed oligonucleotides was then used in a Golden Gate reaction as described above, using BpiI-HF (NEB) as restriction and pBP_sgRNA (Addgene number: 190128) as the target vector, generating the level 0 plasmid pBP_recA_sgRNA or pBP_nonsense_sgRNA respectively. Subsequently, this plasmid was used in a Golden Gate reaction using pMB1-B (Addgene number 190115) and BsaI-HF (NEB) as restriction enzyme, generating pMB1-B-recA_sgRNA. Lastly, pMB1-B-recA was recombined with pMB1-A-pBAD-dCas9 (Addgene number: 190132) in the target plasmid pMB2a-tet (a modified version of Addgene number: 190120 containing TetR instead of KanR) generating the plasmid pMB2a-pBAD_dCas-recA_sgRNA and pMB2a-pBAD_dCas9-nonsense_sgRNA. The plasmids were confirmed via sequencing with the oligonucleotides mentioned in the chapter above. Both plasmids were subsequently transformed into A. baumannii ATCC17978 WT, *ΔrecA*, *ΔumuDAB*, *ΔcsuD* (Moon et al., 2017)(kindly provided by Mario Feldman), and *Δbfmr* (EGA496, kindly provided by Edward Geisinger, Northeastern University) via electroporation.

### Microscopy and GFP reporter measurements

To perform confocal microscopy experiments, we used a Zeiss Axio Observer. Z1/7. The excitation/emission used were 280/618 nm for the red channel, and 488/509 nm for the green channel. We formed biofilms in wells, and after 48 h, 0.5xMIC Cip was added underneath the pellicles and dishes were incubated statically for an additional 48 hrs. Pellicles were collected from mock -treated and Cip-treated wells on a glass coverslip treated with 0.1 mg/mL of polylysine to facilitate attachment. The side of the pellicle in direct contact with the liquid medium was in contact with the glass, and the side of the biofilm in contact with air remained as such. Z-stacks were of 1µm.

For the GFP reporter measurements, GFP fluorescence was measured in a plate reader (Biotek synergy H1) for at least 24 h. at 25°C without shaking. To start the experiment, cultures were diluted 1:100 in YT medium in a 96 multi-well dish. Green fluorescence (Excitation 479 nm and emission 520 nm) and OD_600_ were measured every 15 min. Growth and fluorescence data were gathered at least in triplicate for each strain.

### MIC and Biofilm Eradication Assays, and Live/Dead determination

Standard assays were used to test for MIC (Andrews, 2001) in planktonic shaking cultures. Pellicles were set up as described above and incubated statically at 25°C for the number of hours indicated in the respective figure legends, at which time free cells (not in biofilms) were removed, and wells washed thrice with 1X PBS. YT medium containing Gm, at a range of concentrations shown in the respective figures was added to each washed well. The plate was then incubated statically for 24 hrs. at 25°C, after which the spent medium was removed, and wells were washed three times with PBS. Crystal Violet staining was then carried out as before (Chen et al., 2015). Plating for viable counts was performed as described earlier.

## Acknowledgments

We would like to acknowledge the members of the Godoy, Chai, and Geisinger labs at Northeastern University for helpful comments and discussion. We would like to thank G. Bou from Servicio de Microbiologia, Complejo Hospitalario Universitario, La Coruña, Spain for the Δ*recA* strain, Luis Actis from the Department of Microbiology, Miami University, for the anti-Csu antibody, Stephen Lory from Harvard Medical School for access to the AB7505 transposon collection, Mario Feldman from Washington University St. Louis Medical School for the for Δ*csuD* strain and Edward Geisinger from the Department of Biology, Northeastern University for the *ΔbfmR* strain (EGA496). M.D. and S.R. were funded by the Undergraduate Research and Fellowships office at Northeastern University. V.G.G used funding from a stipend from NUSci, an Inclusive Excellence award form HHMI, Y.C. from an NSF grant (MCB1651732), and A.R. by a Northeastern University Provost Dissertation Completion Fellowship.

## Author Contributions

V.G.G., and C.C. initiated research in discussions with YC. B. I. performed preliminary experiments. C.C., P.M., A.R., M. B., B. N., S. R., M.D., W. F., and V.G.G. performed experiments and analyzed data. C.C. and V.G.G. wrote the manuscript.

## Supplementary data (Table S1 and Figures S1-S4)

**Table S1.**
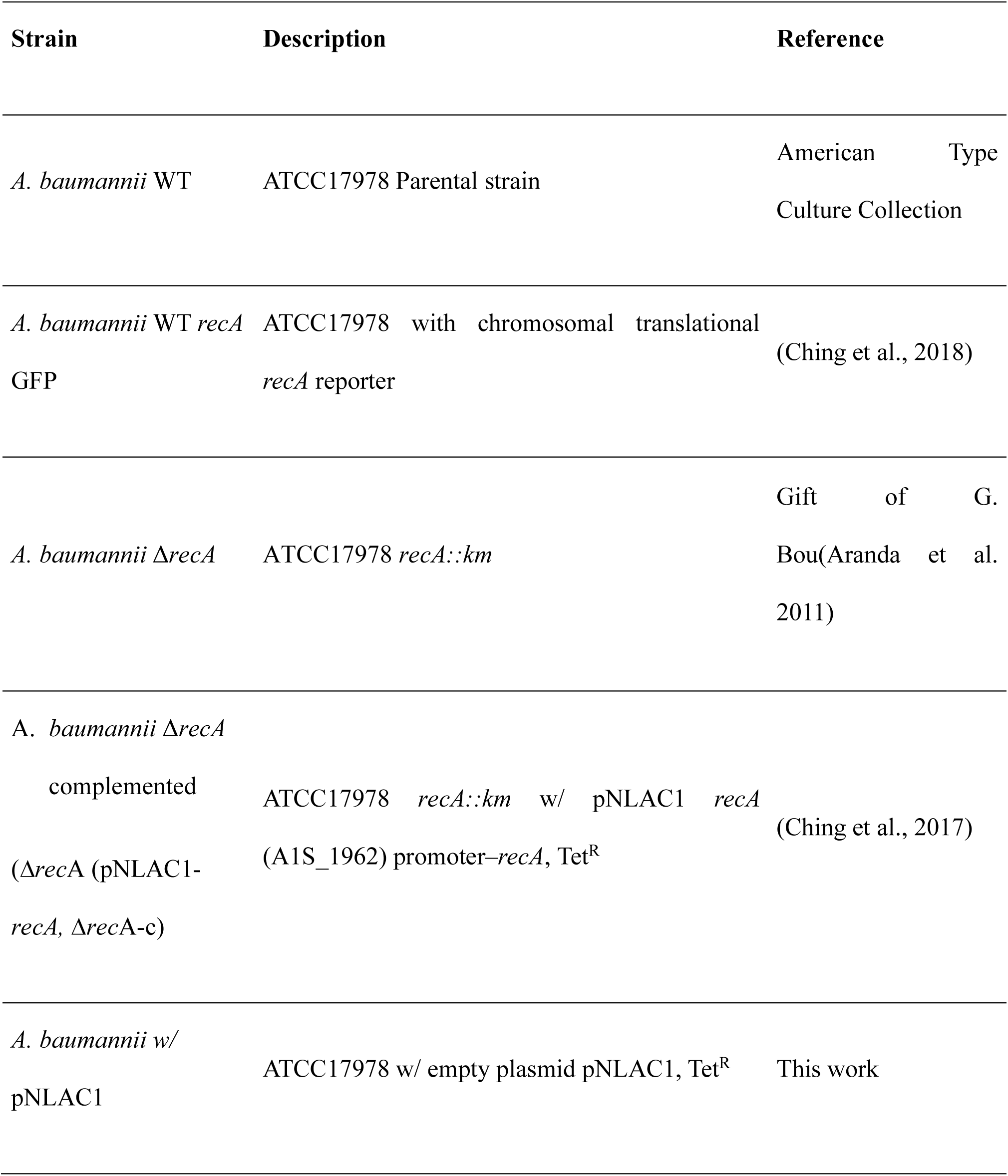
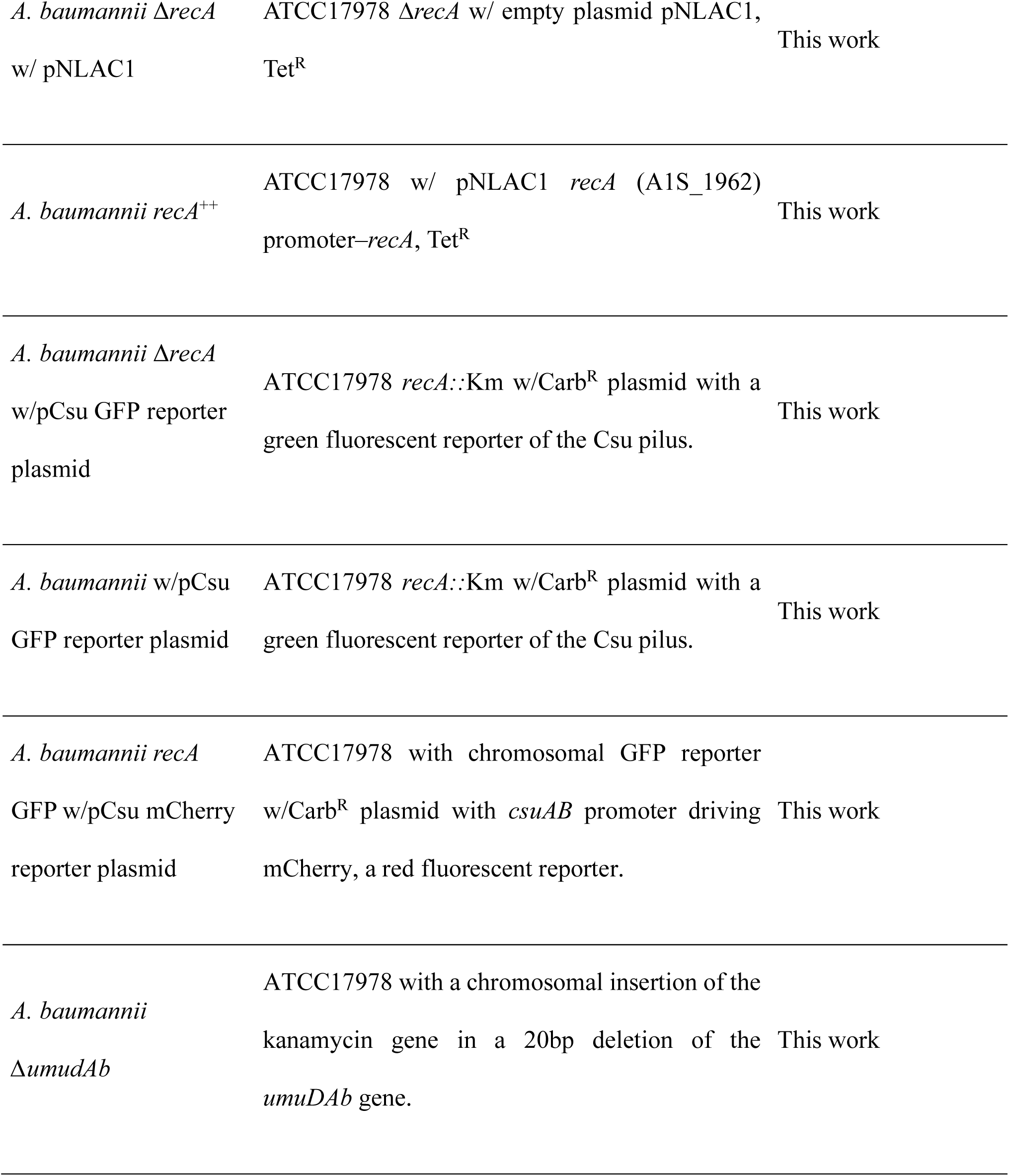
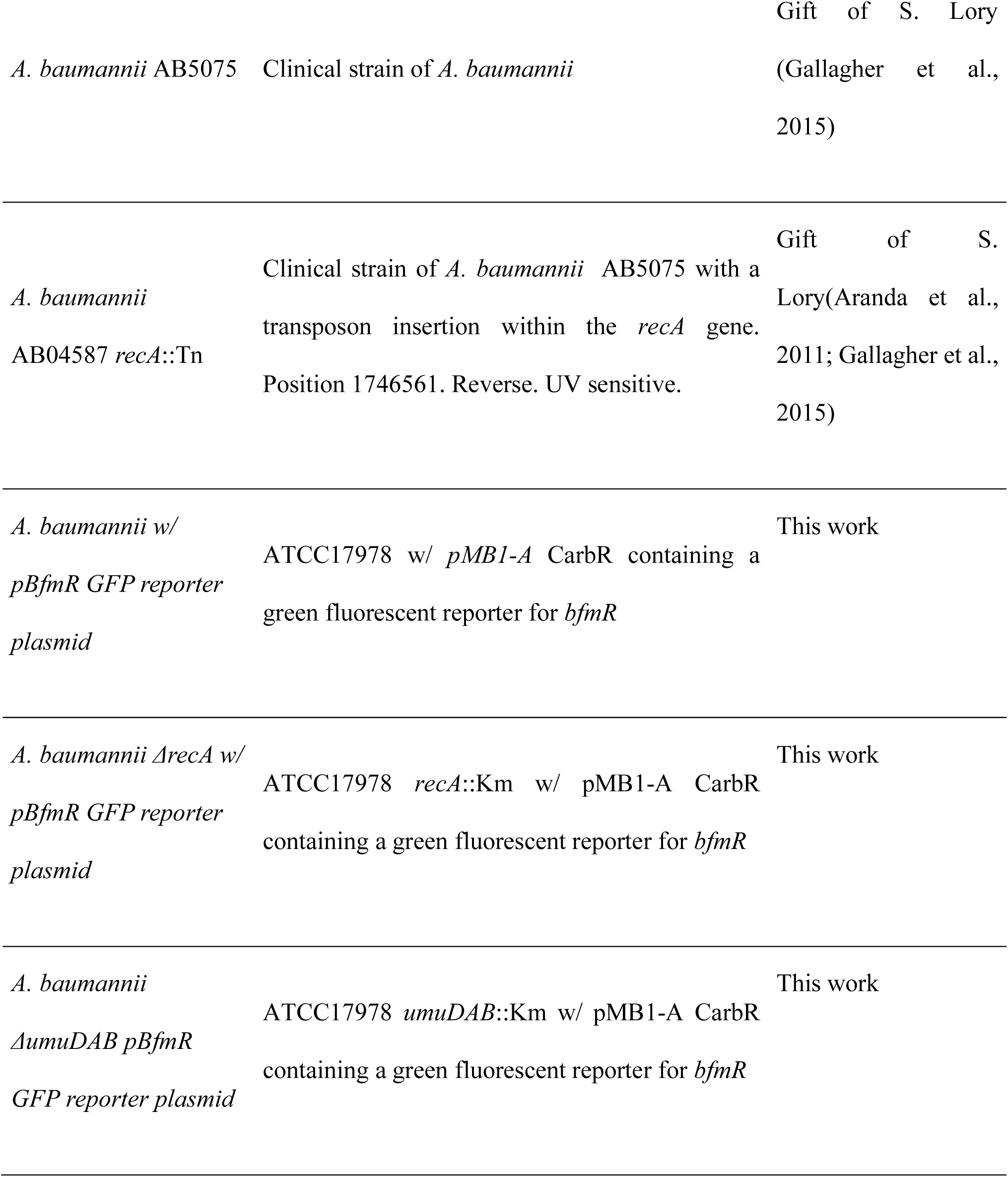
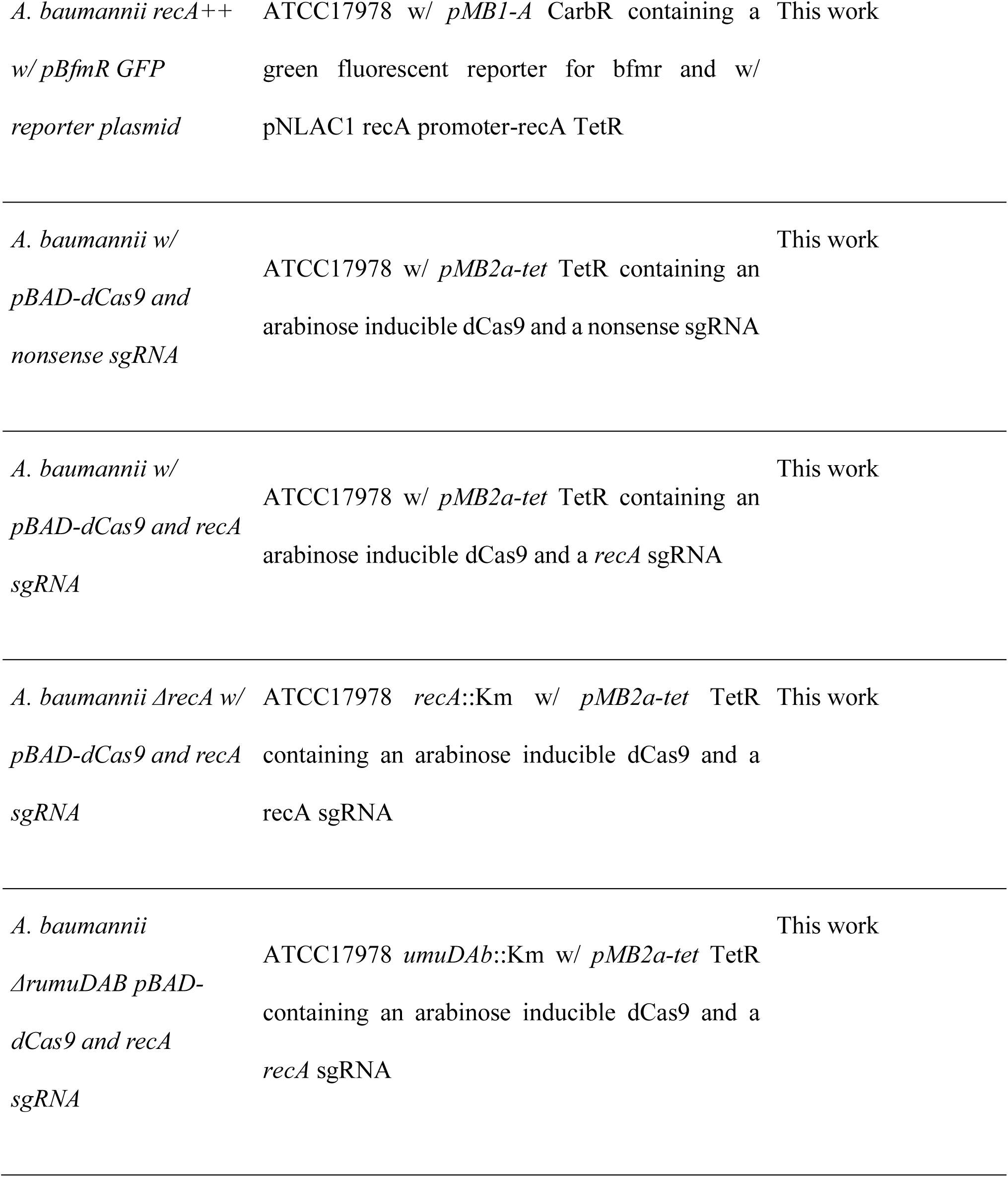
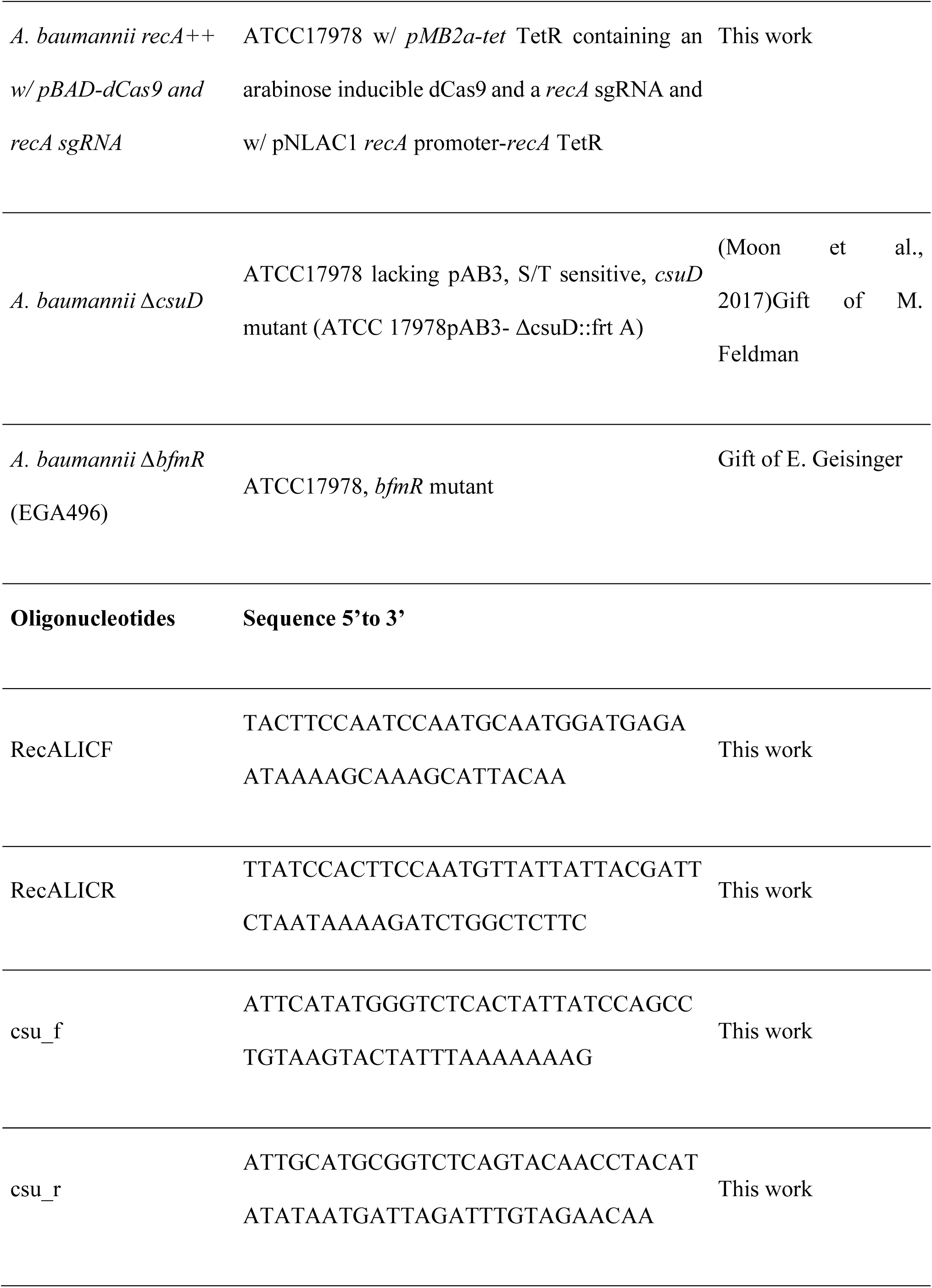
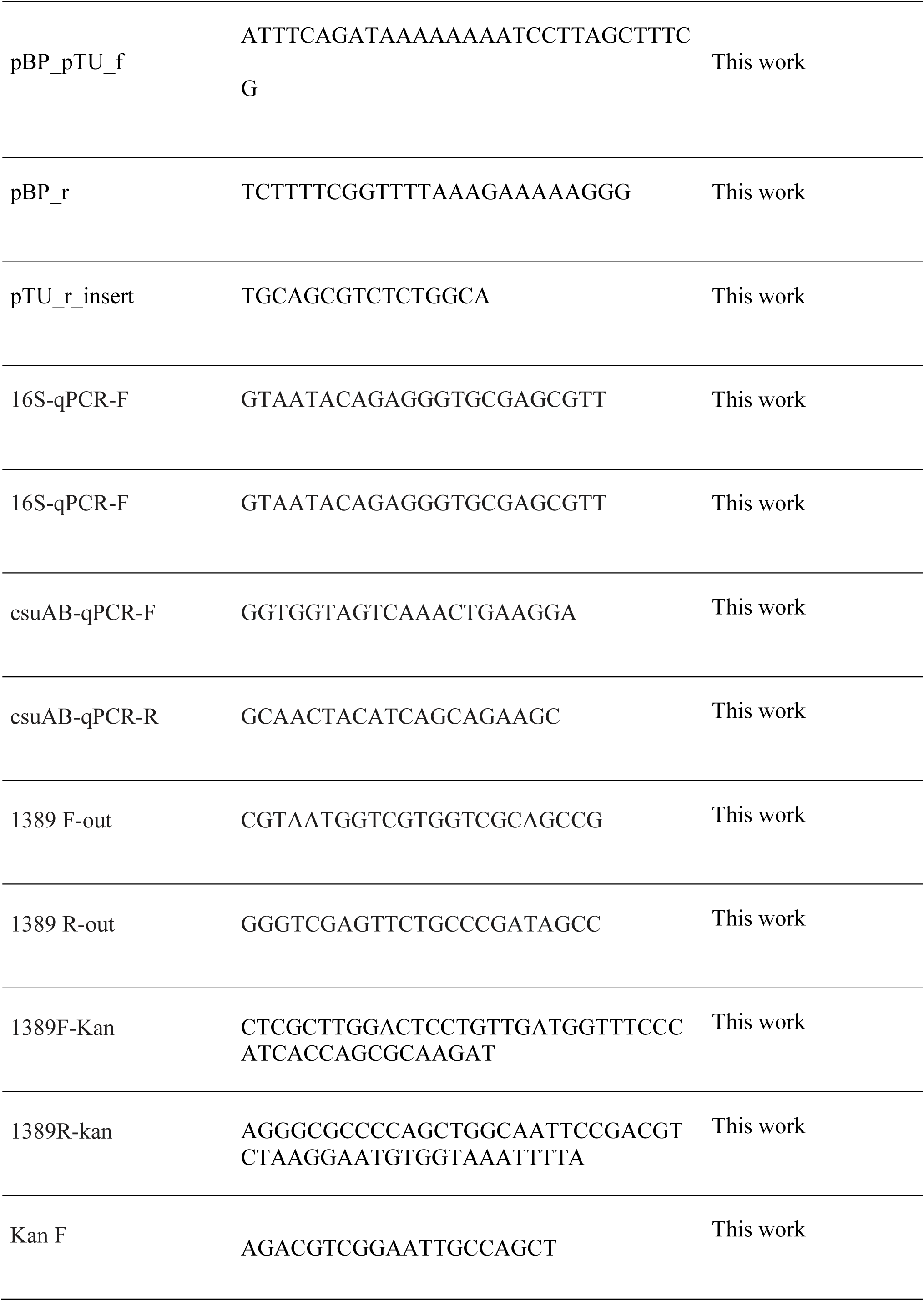
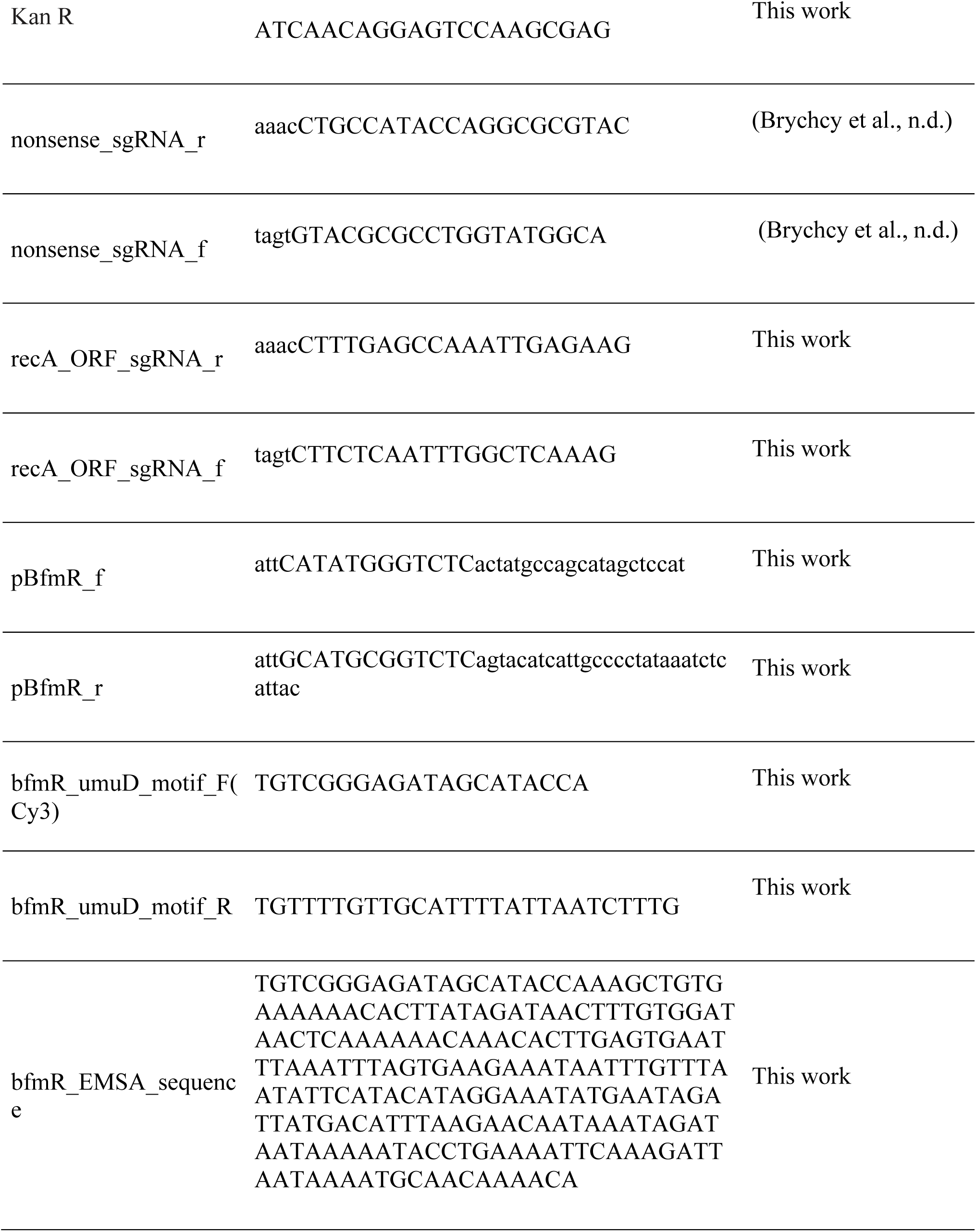
Strains and Oligonucleotides used in this study.

**Figure S1.**
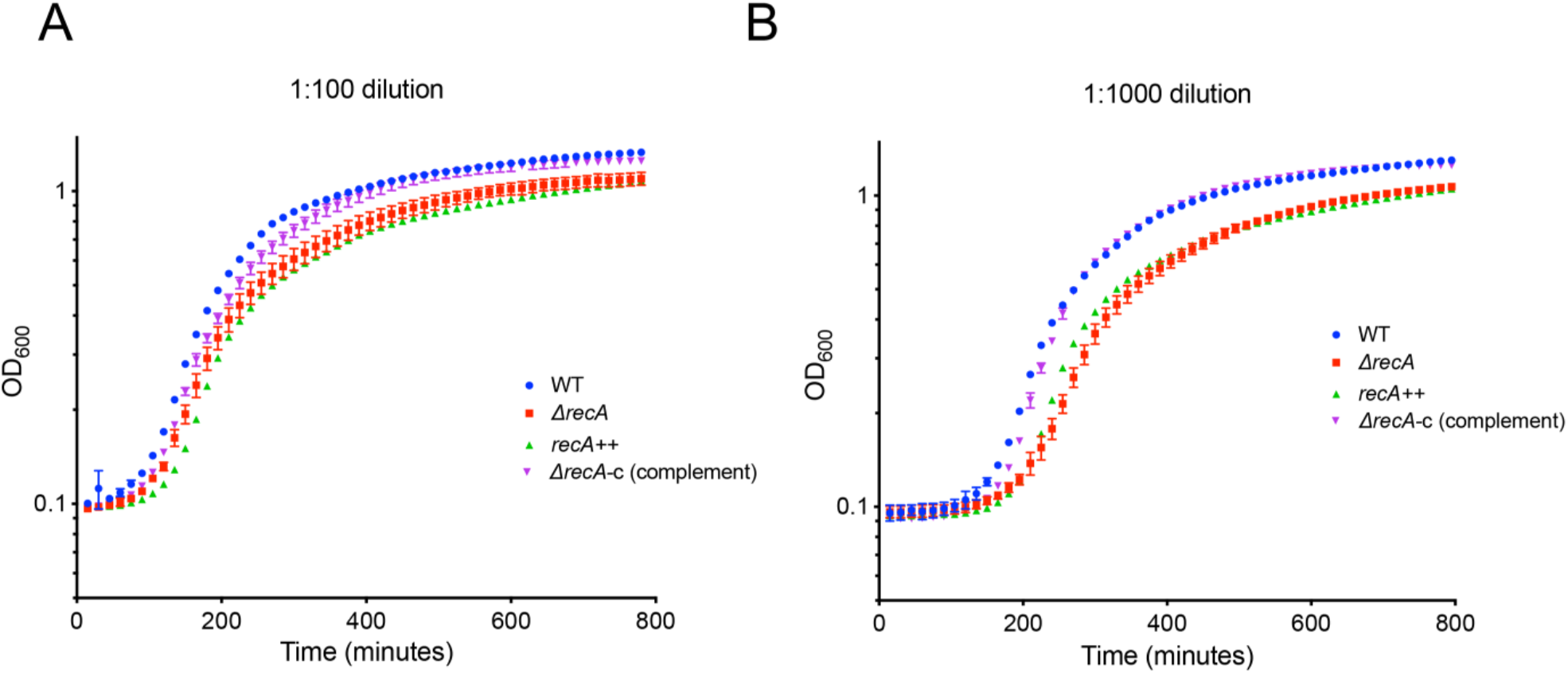
Δ*recA* growth is fully complemented by a plasmid borne copy of *recA* under its own promoter. Cells were diluted at (A) 1:100 and (B) 1:1000 in YT medium and OD600 was measured in a plate reader every 15 mins at 37°C with shaking. Error bars represent standard deviation of the mean of three experiments.

**Figure S2.**
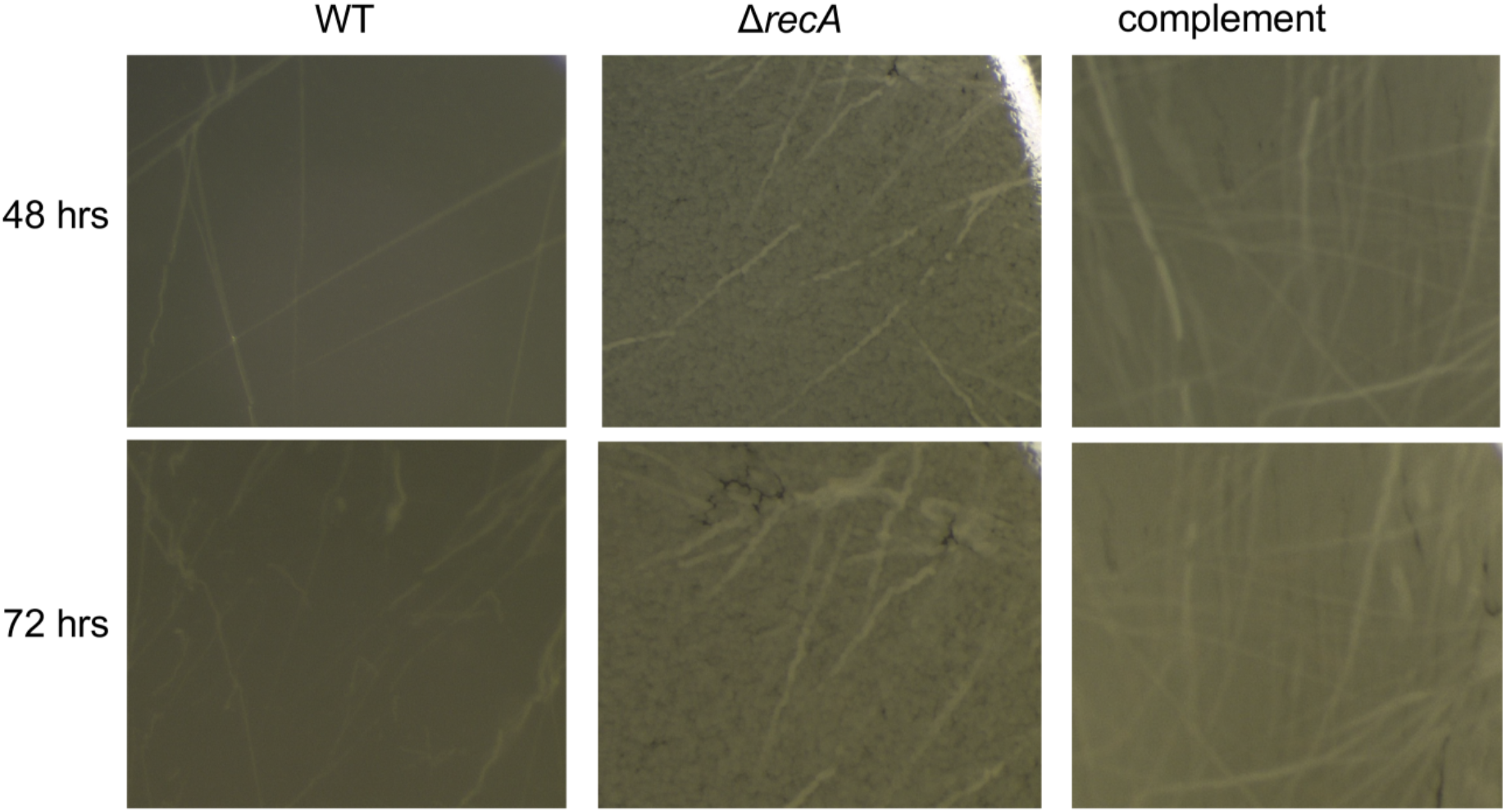
Δ*recA* pellicle biofilms formed at the air-surface interface have different surface features compared to WT. Magnified images of the pellicle surface formed by WT compared to Δ*recA* and the complemented strain at 48 and 72 hours. The WT strain has a smooth surface with striations like complemented strain. The surface of the Δ*recA* biofilm is more granular.

**Figure S3.**
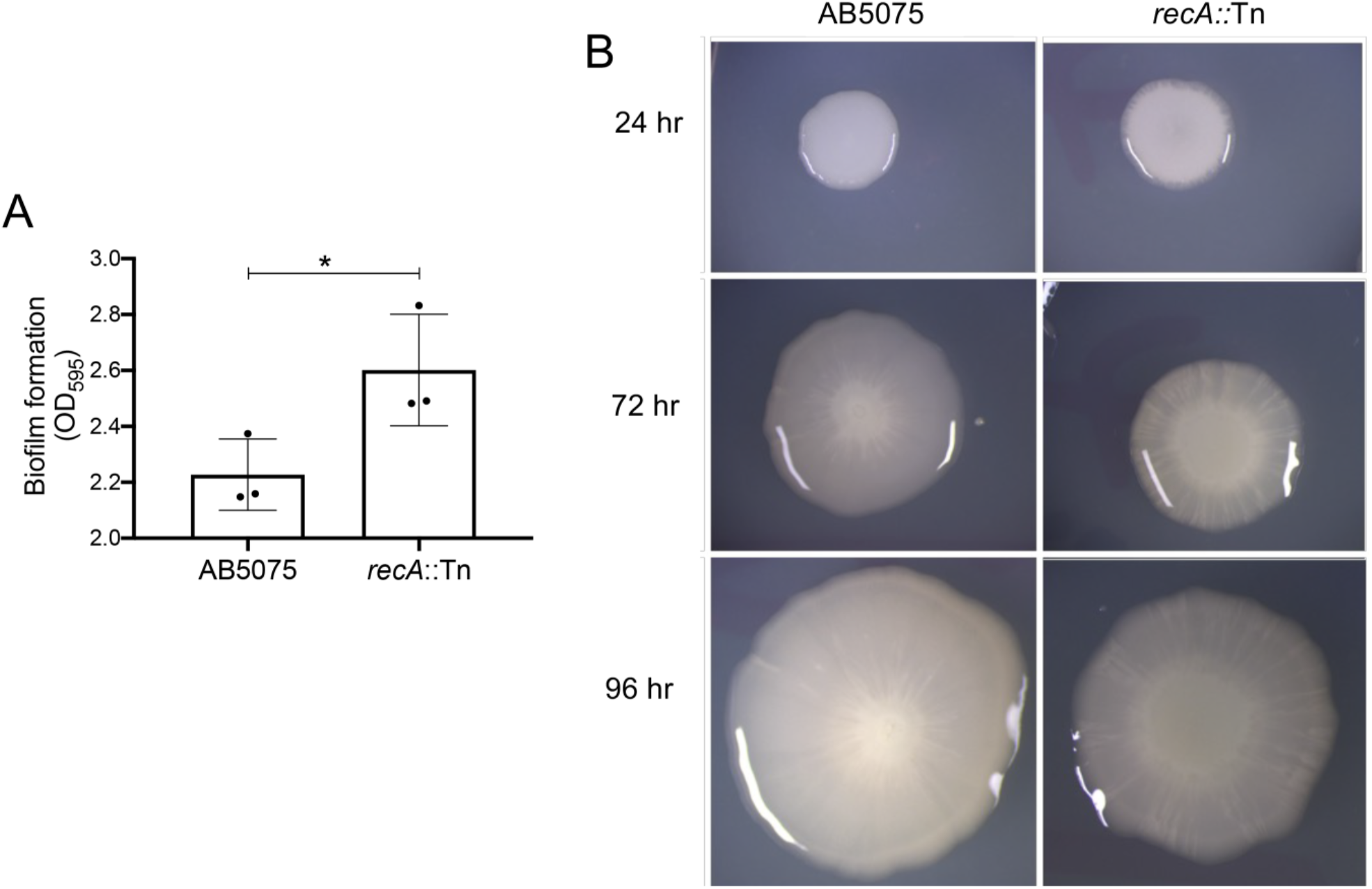
*A. baumannii* AB5075 biofilms are influenced by RecA. **(A)** Absorbance at 595 nm for crystal violet staining, at 48 hrs., of polystyrene-adhered AB5075 and AB5075 *recA*::Tn cells from pellicle biofilms formed in YT medium. Note the Y-axis start at 2.0. An unpaired T-Test was used for statistical analysis relative to the parental strain, *= P< 0.05 **(B)** Colony biofilms at 24, 48 and 72 hrs. in R2 medium with 1.5% agar instead of YT. The colony biofilms of the AB5705 *recA::*Tn have a different morphology and smaller size than the parental strain.

**Figure S4.**
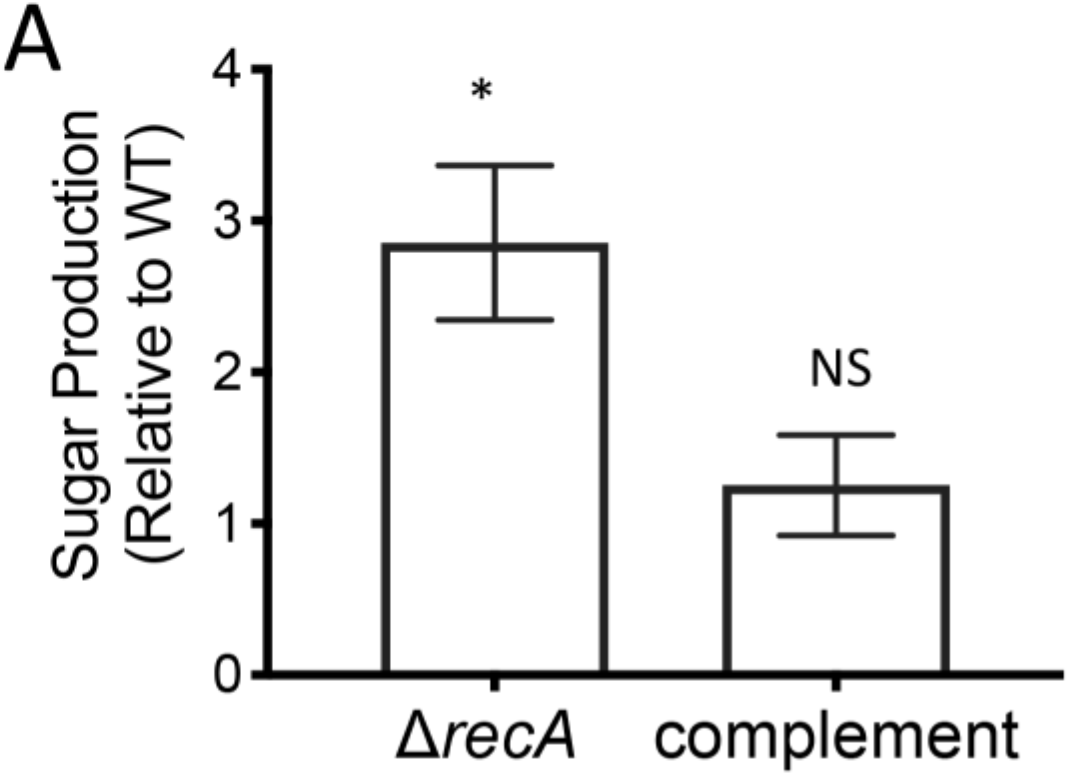
Δ*recA* biofilms have increased observable matrix compared to WT biofilms. (**A**) The matrix fraction isolated from Δ*recA* cells has a shift towards increased total carbohydrate content. Total extracellular matrix was extracted from saturated stationary phase cultures. Total carbohydrates were measured from WT, Δ*recA* and complemented Δ*recA* cells. Experiments were performed in triplicate. Error bars represent standard deviation. An unpaired two-tailed T-Test was used for statistical analysis relative to WT strain; * = P <0.05, NS = Not Significant.

**Figure S5.**
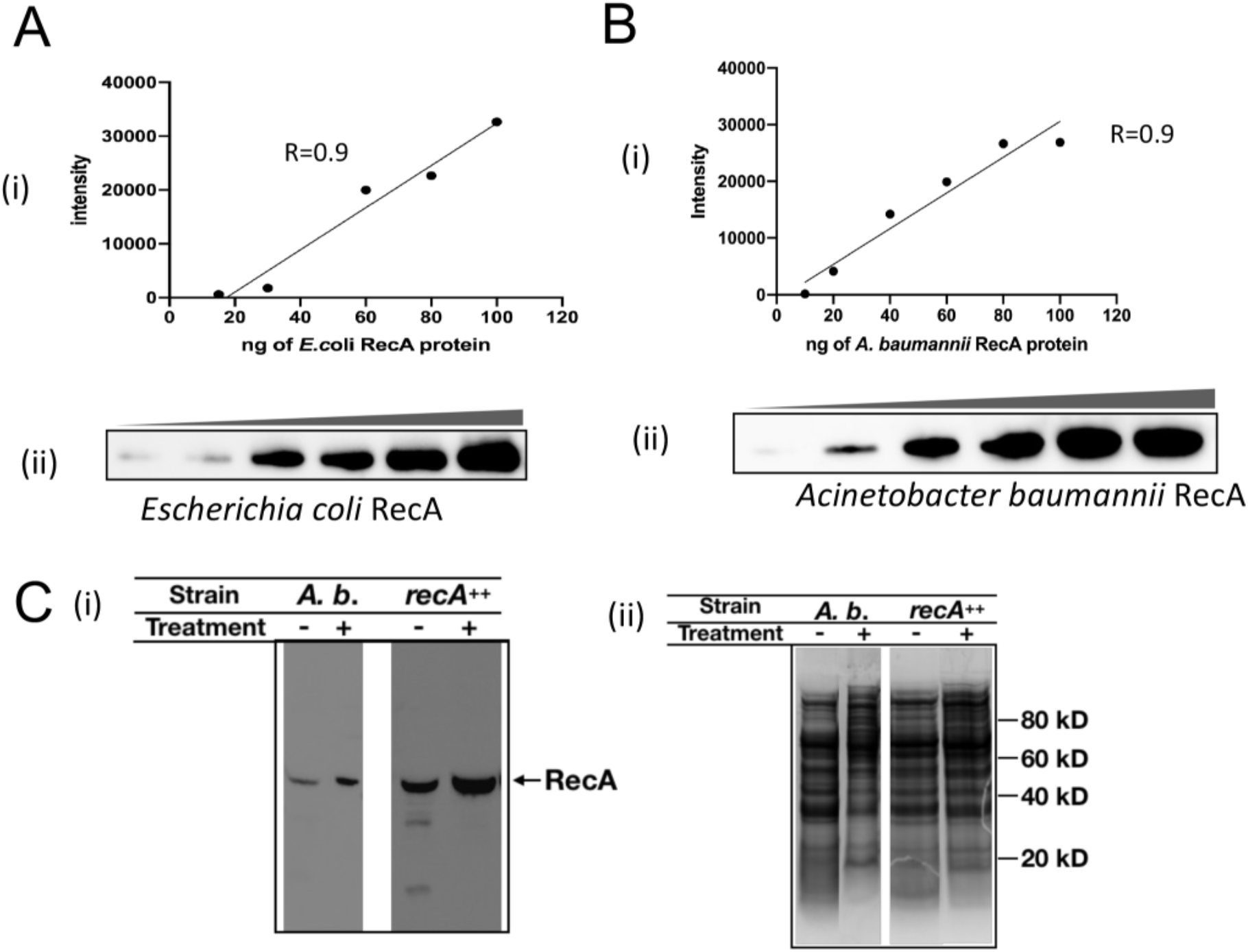
*recA^++^* produces has 3-5-fold more RecA than WT strain. Recombinant *E. coli* RecA an *A. baumannii* RecA were purified as indicated in Materials and Methods. Increasing amounts of either protein were separated by PAGE and an immunoblot was carried out with the same dilution of anti-RecA antibody (Abcam, 1:10,000) and conditions. (A) (i) *E. coli* RecA standard curve (15, 30, 60, 80, 100 and 200 ng) and (B) (ii) *A. baumannii RecA* standard curve (10, 20, 40, 60, 80, and 100 ng). No significant differences in recognition by the antibody were observed. Blot was repeated three times. Representatives immunoblot for each curve (A(ii) and B(ii)) are shown. (C) (i) Immunoblot for RecA of extracts of *A. baumannii* cells with (+) or without (-) 3 hr 10x MIC Cip exposure to induce the DNA damage response. Density analysis with ImageJ of the bands showed that the *recA*^++^ strain has between 3-5-fold more RecA than the WT strain. The samples shown were separated in the same gel and divided electronically for figure clarity. (ii) Separation of cell free extracts of the respective samples in (i) by SDS-PAGE on a 12% Bis-Tris gel in MOPS buffer to demonstrate that similar total protein was loaded per well. Estimated molecular weight sizes are shown in the right-hand size of the gel. The calculations are the result of 4 biological replicates.

**Figure S6.**
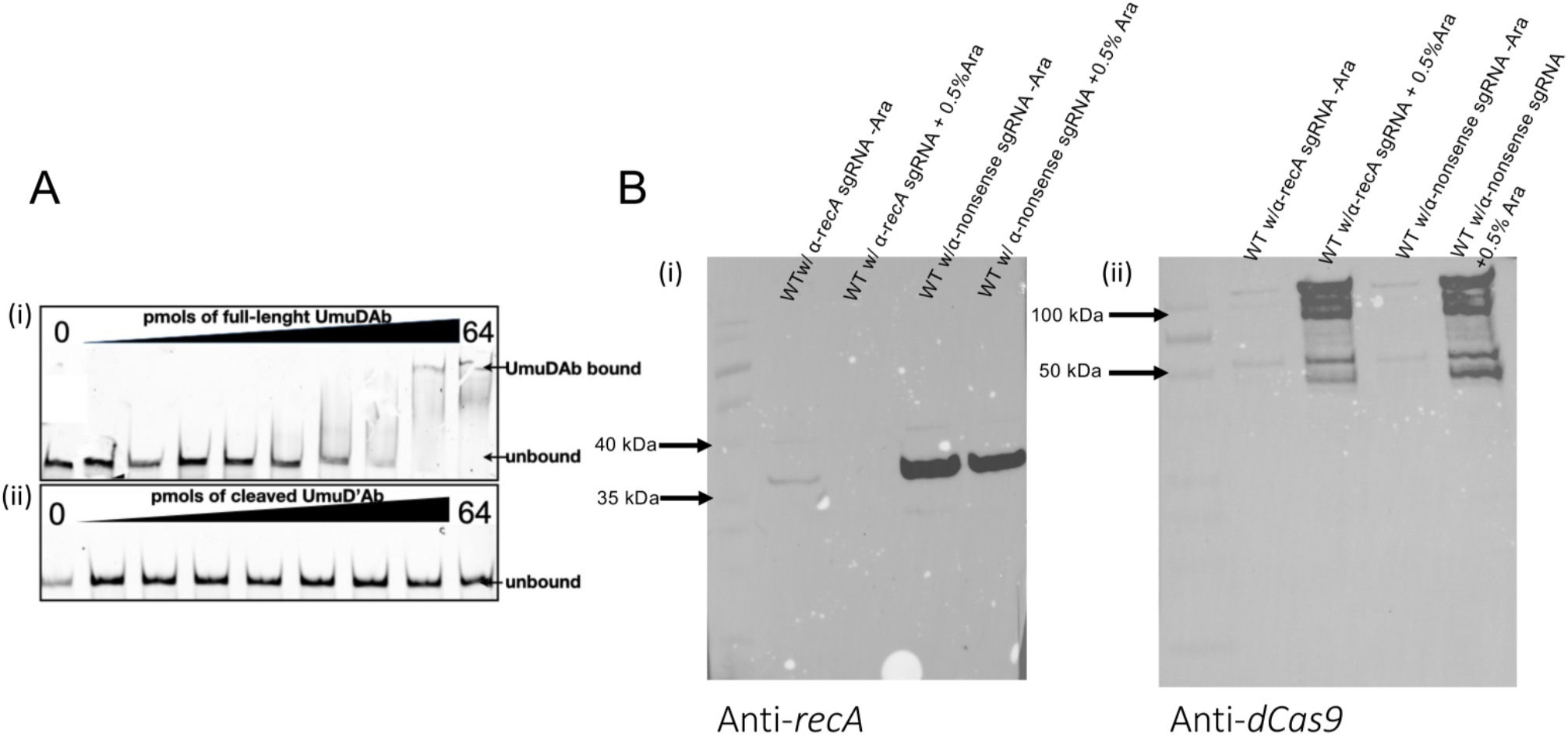
(A) Electrophoresis mobility shift assay (EMSA) of purified (i) full length UmudAb and (ii) cleaved UmuDAb to sequences containing the *bfmrR* promoter. The recombinant proteins were purified in *E. coli* from his-tagged versions of full length and cleaved UmuDAb. B) The plasmid-based CRSPRi we developed inhibits *recA* expression only in cells containing the sgRNA targeting the *recA* gene. Immunoblot for (i) RecA and (ii) dCas9 in CRISPRi knock down strains of WT background with *recA* and nonsense sgRNA with (+) or without (-) 0.5% Arabinose.

## References

1. Alam K, Alhhazmi A, Decoteau JF, Luo Y, Geyer CR, Alam K, Alhhazmi A, Decoteau JF, Luo Y, Geyer CR. 2016. RecA Inhibitors Potentiate Antibiotic Activity and Block Evolution of Antibiotic Resistance Article RecA Inhibitors Potentiate Antibiotic Activity and Block Evolution of Antibiotic Resistance. Cell Chem Biol 23:381–391. doi:10.1016/j.chembiol.2016.02.010

2. Anderl JN, Franklin MJ. 2000. Role of Antibiotic Penetration Limitation in Klebsiella pneumoniae Biofilm Resistance to Ampicillin and Ciprofloxacin. Antimicrob Agents Chemother 44:1818– 1824.

3. Andrews JM. 2001. Determination of minimum inhibitory concentrations. Journal of Antimicrobial Chemotherapy 5–16.

4. Aranda J, Bardina C, Beceiro A, Rumbo S, Cabral MP, Barbé J, Bou G. 2011. Acinetobacter baumannii RecA protein in repair of DNA damage, antimicrobial resistance, general stress response, and virulence. J Bacteriol 193:3740–7. doi:10.1128/JB.00389-11

5. Asally M, Kittisopikul M, Rue P, Du Y, Hu Z, Cagatay T, Robinson AB, Lu H, Garcia-Ojalvo J, Suel GM. 2012. Localized cell death focuses mechanical forces during 3D patterning in a biofilm.

6. Proceedings of the National Academy of Sciences 109:18891–18896. doi:10.1073/pnas.1212429109

7. Bales PM, Renke EM, May SL, Shen Y, Nelson DC. 2013. Purification and Characterization of Biofilm-Associated EPS Exopolysaccharides from ESKAPE Organisms and Other Pathogens. PLoS One 8. doi:10.1371/journal.pone.0067950

8. Blair JMA, Webber MA, Baylay AJ, Ogbolu DO, Piddock LJ v. 2014. Molecular mechanisms of antibiotic resistance. Nat Rev Microbiol 13:42–51. doi:10.1038/nrmicro3380

9. Brossard KA, Campagnari AA. 2012. The Acinetobacter baumannii Biofilm-Associated Protein Plays a Role in Adherence to Human Epithelial Cells. Infect Immun 228–233. doi:10.1128/IAI.05913-11

10. Brychcy M, Kokodynski A, Lloyd D, Godoy VG. n.d. AspFlex: molecular tools to study gene expression and regulation in. bioRxiv. doi:10.1101/2023.03.17.533205

11. Cafarelli TM, Rands TJ, Benson RW, Rudnicki PA, Lin I, Godoy VG. 2013. A single residue unique to dinb-like proteins limits formation of the polymerase IV multiprotein complex in Escherichia coli. J Bacteriol 195:1179–1193. doi:10.1128/JB.01349-12

12. Cafarelli TM, Rands TJ, Godoy VG. 2014. The DinB.RecA Complex of Escherichia coli Mediates an Efficient and High-Fidelity Response to Ubiquitous Alkylation Lesions. Mutat Res Genet Toxicol Environ Mutagen 51:229–235. doi:10.1002/em

13. Chen Y, Gozzi K, Yan F, Chai Y. 2015. Acetic acid acts as a volatile signal to stimulate bacterial biofilm formation. mBio 6:1–13. doi:10.1128/mBio.00392-15

14. Ching C, Gozzi K, Heinemann B, Chai Y, Godoy VG. 2017. RNA-Mediated cis Regulation in Acinetobacter baumannii Modulates Stress-Induced Phenotypic Variation. J Bacteriol 199:1–15.

15. Ching C, Yang B, Onwubueke C, Lazinski D, Camilli A, Godoy VG. 2018. Lon protease has multifaceted biological functions in *Acinetobacter baumannii*. J Bacteriol 201:1–12. doi:10.1128/JB.00536-18

16. Choi AH, Slamti L, Avci FY, Pier GB. 2009. The pgaABCD Locus of Acinetobacter baumannii Encodes the Production of Poly-␤-1-6-N-Acetylglucosamine, Which Is Critical for Biofilm Formation. J Bacteriol 191:5953–5963. doi:10.1128/JB.00647-09

17. Cirz RT, Jones MB, Gingles NA, Minogue TD, Jarrahi B, Peterson SN, Romesberg FE. 2007. Complete and SOS-mediated response of Staphylococcus aureus to the Antibiotic Ciprofloxacin. J Bacteriol 189:531–539. doi:10.1128/JB.01464-06

18. Costa SB, Campos ACC, Pereira ACM, de Mattos-Guaraldi AL, Junior RH, Rosa ACP, Asad LMBO. 2014. Adherence to abiotic surface induces SOS response in Escherichia coli K-12 strains under aerobic and anaerobic conditions. Microbiology (N Y) 160:1964–1973. doi:10.1099/mic.0.075317-0

19. Costerton JW, Lewandowski Z, Caldwell DE, Korber DR, Sn S, Lappin-scott HM. 1995. Microbial biofilms. Annu Rev Microbiol 711–745.

20. Cox JM, Li H, Wood EA, Chitteni-Pattu S, Inman RB, Cox MM. 2008. Defective dissociation of a “slow” RecA mutant protein imparts an Escherichia coli growth defect. Journal of Biological Chemistry 283:24909–24921. doi:10.1074/jbc.M803934200

21. Draughn GL, Milton ME, Feldmann EA, Bobay BG, Roth BM, Olson AL, Thompson RJ, Actis LA, Davies C, Cavanagh J. 2018. The Structure of the Biofilm-controlling Response Regulator BfmR from Acinetobacter baumannii Reveals Details of Its DNA-binding Mechanism. J Mol Biol 430:806–821. doi:10.1016/j.jmb.2018.02.002

22. Ducret A, Quardokus E, Brun Y. 2016. MicrobeJ, a high throughput tool for quantitative bacterial cell detection and analysis. Nature Microbiol 1:1–7. doi:10.1038/nmicrobiol.2016.77

23. Erlandsen SL, Kristich CJ, Dunny GM, Wells CL. 2004. High-resolution Visualization of the Microbial Glycocalyx with Low-voltage Scanning Electron Microscopy : Dependence on Cationic Dyes The Journal of Histochemistry & Cytochemistry. Journal of Histochemistry and Cytochemistry 52:1427–1435. doi:10.1369/jhc.4A6428.2004

24. Eze EC, Chenia HY, el Zowalaty ME. 2018. Acinetobacter baumannii biofilms : effects of physicochemical factors, virulence, antibiotic resistance determinants, gene regulation, and future antimicrobial treatments. Infect Drug Resist 11:2277–2299.

25. Farrow JM, Wells G, Pesci EC. 2018. Desiccation tolerance in Acinetobacter baumannii is mediated by the two-component response regulator BfmR. PLoS One 13:1–25. doi:10.1371/journal.pone.0205638

26. Fuchs RP, Fujii S. 2013. Translesion DNA Synthesis and Mutagenesis in Prokaryotes. Cold Spring Harb Perspect Biol 1–22.

27. Gaddy JA, Actis LA. 2009. Regulation of Acinetobacter baumannii biofilm formation. Future Microbiol 4:273–278. doi:10.2217/fmb.09.5.Regulation

28. Gallagher LA, Ramage E, Weiss EJ, Radey M, Hayden HS, Held KG, Huse HK, Zurawski D v., Brittnacher MJ, Manoil C. 2015. Resources for genetic and genomic analysis of emerging pathogen Acinetobacter baumannii. J Bacteriol 197:2027–2035. doi:10.1128/JB.00131-15

29. Geisinger E, Isberg RR. 2015. Antibiotic Modulation of Capsular Exopolysaccharide and Virulence in Acinetobacter baumannii. PLoS Pathog 11:1–27. doi:10.1371/journal.ppat.1004691

30. Ghodke H, Paudel BP, Lewis JS, Jergic S, Gopal K, Romero ZJ, Wood EA, Woodgate R, Cox MM, Oijen AMV. 2019. Spatial and temporal organization of reca in the escherichia coli dna-damage response. Elife 8:1–37. doi:10.7554/eLife.42761

31. Gozzi K, Ching C, Paruthiyil S, Zhao Y, Godoy VG, Chai Y. 2017. Bacillus subtilis utilizes the DNA damage response to manage multicellular development. NPJ Biofilms Microbiomes 3:8. doi:10.1038/s41522-017-0016-3

32. Gradia S, Ishida JP, Tsai M, Jeans C, Tainer JA, States U, Bioimaging I, Berkeley L, States U, Berkeley L, States U, States U, States U. 2017. MacroBac: New technologies for robust and efficient large-scale production of recombinant multi-protein complexes. Methods Enzymol. doi:10.1016/bs.mie.2017.03.008.MacroBac

33. Hardouin J, Chabane YN, Marti S, Rihouey C. 2014. Characterisation of Pellicles Formed by Acinetobacter baumannii at the Air-Liquid Interface. PLoS One 9. doi:10.1371/journal.pone.0111660

34. Hare JM, Adhikari S, Lambert K V, Hare AE, Alison N. 2013. The Acinetobacter regulatory UmuDAb protein cleaves in response to DNA damage with chimeric LexA/UmuD characteristics. FEMS Microbiol Lett 334:57–65. doi:10.1111/j.1574-6968.2012.02618.x.The

35. Hare JM, Ferrell JC, Witkowski T a, Grice AN. 2014. Prophage induction and differential RecA and UmuDAb transcriptome regulation in the DNA damage responses of Acinetobacter baumannii and Acinetobacter baylyi. PLoS One 9:e93861. doi:10.1371/journal.pone.0093861

36. He X, Lu F, Yuan F, Jiang D, Zhao P, Zhu J, Chene H, Cao J, Lu G. 2015. Biofilm Formation Caused by Clinical Acinetobacter baumannii Isolates Is Associated with Overexpression of the AdeFGH Efflux Pump. Antimicrob Agents Chemother 59:4817–4825. doi:10.1128/AAC.00877-15

37. Inagaki S, Fujita K, Nagayama K, Funao J, Effects OT. 2009. Effects of recombinase A deficiency on biofilm formation by Streptococcus mutans. Oral Microbiol Immunol 104–108.

38. Jacobs AC, Thompson MG, Black CC. 2014. AB5075, a Highly Virulent Isolate of Acinetobacter baumannii, as a. mBio 5:1–10. doi:10.1128/mBio.01076-14.Editor

39. Jiao Y, Cody GD, Harding AK, Wilmes P, Schrenk M, Wheeler KE, Banfield JF, Thelen MP, Carolina N. 2010. Characterization of Extracellular Polymeric Substances from Acidophilic Microbial Biofilms 1 †. Appl Environ Microbiol 76:2916–2922. doi:10.1128/AEM.02289-09

40. Kreuzer KN. 2013. DNA Damage Responses in Prokaryotes: Regulating Gene Expression, Modulating Growth Patterns, and Manipulating Replication Forks. Cold Spring Harb Perspect Biol 1–23.

41. Linares JF, Gustafsson I, Baquero F, Martinez JL. 2006. Antibiotics as intermicrobial signaling agents instead of weapons. Proc Natl Acad Sci U S A 103:19484–19489. doi:10.1073/pnas.0608949103

42. Little JW, Mount DW. 1982. The SOS Regulatory Escherichia coli System of Review 29:1982.

43. Loehfelm TW, Luke NR, Campagnari AA. 2008. Identification and characterization of an Acinetobacter baumannii biofilm-associated protein. J Bacteriol 190:1036–1044. doi:10.1128/JB.01416-07

44. Luke NR, Sauberan SL, Russo TA, Beanan JM, Olson R, Loehfelm TW, Cox AD, Michael FS, Vinogradov E v, Campagnari AA. 2010. Identification and Characterization of a Glycosyltransferase Involved in Acinetobacter baumannii Lipopolysaccharide Core Biosynthesis 1. Infect Immun 78:2017–2023. doi:10.1128/IAI.00016-10

45. MacGuire AE, Ching MC, Diamond BH, Kazakov A, Novichkov P, Godoy VG. 2014. Activation of phenotypic subpopulations in response to ciprofloxacin treatment in Acinetobacter baumannii. Mol Microbiol 92:138–52. doi:10.1111/mmi.12541

46. Moon KH, Weber BS, Feldman F. 2017. Subinhibitory Concentrations of Trimethoprim and Sulfamethoxazole Acinetobacter baumannii through Inhibition of Csu Pilus Expression. Antimicrob Agents Chemother 61:1–18.

47. Moore SJ, Lai HE, Kelwick RJR, Chee SM, Bell DJ, Polizzi KM, Freemont PS. 2016. EcoFlex: A Multifunctional MoClo Kit for E. coli Synthetic Biology. ACS Synth Biol 5:1059–1069. doi:10.1021/acssynbio.6b00031

48. Nguyen LH, Jensen DB, Burgess RR. 1993. Overproduction and purification of σ32, the Escherichia coli heat shock transcription factor. Protein Expr Purif 4:425–433.

49. Norton MD, Spilkia AJ, Godoy VG. 2013. Antibiotic Resistance Acquired through a DNA Damage-Inducible Response in Acinetobacter baumannii. J Bacteriol 195:1335–45. doi:10.1128/JB.02176-12

50. Pakharukova N, Tuittila M, Paavilainen S, Malmi H, Parilova O, Teneberg S, Knight SD, Zavialov A v. 2018. Structural basis for Acinetobacter baumannii biofilm formation. Proc Natl Acad Sci U S A 115:5558–5563. doi:10.1073/pnas.1800961115

51. Peleg AY, Seifert H, Paterson DL. 2008. Acinetobacter baumannii: Emergence of a successful pathogen. Clin Microbiol Rev 21:538–582. doi:10.1128/CMR.00058-07

52. Reasoner DJ, Geldreich EE. 1985. A new medium for the enumeration and subculture of bacteria from potable water. Appl Environ Microbiol 49:1–7. doi:10.1128/aem.49.1.1-7.1985

53. Rice LB. 2008. Federal Funding for the Study of Antimicrobial Resistance in Nosocomial Pathogens : No ESKAPE. J Infect Dis 197:1079–1081. doi:10.1086/533452

54. Rumbo-Feal S, Gómez MJ, Gayoso C, Álvarez-Fraga L, Cabral MP, Aransay AM, Rodríguez-Ezpeleta N, Fullaondo A, Valle J, Tomás M, Bou G, Poza M. 2013. Whole Transcriptome Analysis of Acinetobacter baumannii Assessed by RNA-Sequencing Reveals Different mRNA Expression Profiles in Biofilm Compared to Planktonic Cells. PLoS One 8:1–19. doi:10.1371/journal.pone.0072968

55. Russo TA, Manohar A, Beanan JM, Olson R, MacDonald U, Graham J, Umland TC. 2016. The Response Regulator BfmR Is a Potential Drug Target for Acinetobacter baumannii. mSphere 1:1–19. doi:10.1128/msphere.00082-16

56. Sahu PK, Iyer PS, Oak AM, Pardesi KR, Chopade BA. 2012. Characterization of eDNA from the Clinical Strain Acinetobacter baumannii AIIMS 7 and Its Role in Biofilm Formation. The Scientific World Journal 2012. doi:10.1100/2012/973436

57. Santos-Lopez A, Marshall CW, Scribner MR, Snyder DJ, Cooper VS. 2019. Evolutionary pathways to antibiotic resistance are dependent upon environmental structure and bacterial lifestyle. Elife 8:1–23. doi:10.7554/elife.47612

58. Shaner NC, Steinbach PA, Tsien RY. 2005. A guide to choosing fluorescent proteins. Nat Methods 2:905–909. doi:10.1038/nmeth819

59. Singh R, Sahore S, Kaur P, Rani A, Ray P. 2016. Penetration barrier contributes to bacterial biofilm-associated resistance against only select antibiotics, and exhibits genus-, strain- and antibiotic-specific differences. Pathog Dis 2–7. doi:10.1093/femspd/ftw056

60. Smith MG, Gianoulis T a, Pukatzki S, Mekalanos JJ, Ornston LN, Gerstein M, Snyder M. 2007. New insights into Acinetobacter baumannii pathogenesis revealed by high-density pyrosequencing and transposon mutagenesis. Genes Dev 21:601–14. doi:10.1101/gad.1510307

61. Studier FW. 2005. Protein production by auto-induction in high density shaking cultures. Protein Expr Purif 41:207–234. doi:10.1016/j.pep.2005.01.016

62. Surgalla MJ, Beesley ED. 1969. Congo red-agar plating medium for detecting pigmentation in Pasteurella pestis. Appl Microbiol 18:834–837.

63. Takajashi A, Yomoda S, Ushijima I, Inoue M. 1995. Orfloxacin, norfloxacin and ceftazimide increase the production of alginate and promote the formation of biofilm of Pseudomonase aeruginosa in vitro. Journal of Antimicrobial Chemotherapy.

64. Tashjian TF, Lin I, Belt V, Cafarelli TM, Godoy VG. 2017. RNA primer extension hinders DNA synthesis by Escherichia coli mutagenic DNA polymerase IV. Front Microbiol 8:1–9. doi:10.3389/fmicb.2017.00288

65. Tomaras AP, Dorsey CW, Edelmann RE, Actis LA. 2003. Attachment to and biofilm formation on abiotic surfaces by Acinetobacter baumannii: Involvement of a novel chaperone-usher pili assembly system. Microbiology (N Y) 149:3473–3484. doi:10.1099/mic.0.26541-0

66. Tomaras AP, Flagler MJ, Dorsey CW, Gaddy JA, Actis LA. 2008. Characterization of a two-component regulatory system from Acinetobacter baumannii that controls biofilm formation and cellular morphology. Microbiology (N Y) 154:3398–3409. doi:10.1099/mic.0.2008/019471-0

67. Vlamakis H, Chai Y, Beauregard P, Losick R, Kolter R. 2013. Sticking together: Building a biofilm the Bacillus subtilis way. Nat Rev Microbiol 11:157–168. doi:10.1038/nrmicro2960

68. Walter BM, Cartman ST, Minton NP, Butala M, Rupnik M, B.M. W, S.T. C, N.P. M, M. B, M. R, Walter BM, Cartman ST, Minton NP, Butala M, Rupnik M. 2015. The SOS Response Master Regulator LexA Is Associated with Sporulation, Motility and Biofilm Formation in Clostridium difficile. PLoS One 10:e0144763–e0144763. doi:10.1371/journal.pone.0144763

69. Wang Y, Huang T, Yang Y, Kuo S, Chen C. 2018. Biofilm formation is not associated with worse outcome in Acinetobacter baumannii bacteraemic pneumonia. Sci Rep 1–10. doi:10.1038/s41598-018-25661-9

70. Werner E, Roe F, Bugnicourt A, Franklin MJ, Heydorn A, Molin S, Pitts B, Stewart PS. 2004. Stratified Growth in Pseudomonas aeruginosa Biofilms. Appl Environ Microbiol 70:6188–6196. doi:10.1128/AEM.70.10.6188

71. Witkowski TA, Grice AN, Stinnett DB, Wells WK, Peterson MA, Hare JM. 2016. UmuDAb : An Error-Prone Polymerase Accessory Homolog Whose N-Terminal Domain Is Required for Repression of DNA Damage Inducible Gene Expression in Acinetobacter baylyi. PLoS One 1–19. doi:10.1371/journal.pone.0152013

